# Reorganization of large-scale network dynamics after early visual deprivation

**DOI:** 10.1101/2024.10.11.617823

**Authors:** Caterina A. Pedersini, Bas Rokers, Olivier Collignon

**Author notes:** Corresponding author: Caterina Annalaura Pedersini. **Data, Materials, and Software Availability** The statistical outputs as well as code used to estimate the HMMs, compare the model orders and run the statistical analyses have been deposited in the Open Science Framework (https://osf.io/8a7ft/). **Declaration of Competing Interests** No conflicts of interest, financial or otherwise, are declared by the author(s). **Acknowledgements** We wish to extend our gratitude to Marco Barillari for his help arranging data sharing and providing details about the acquisition protocols. We are also extremely thankful to our blind participants for their invaluable contributions. OC is a senior researcher at FRS-FNRS.

## Abstract

How sensory experience shapes the organization of the human brain connectome remains a central question in neuroscience. Previous studies of congenital blindness have primarily relied on stationary functional connectivity, potentially overlooking how sensory deprivation affects the temporal reconfiguration of large-scale brain networks. Here, we investigated dynamic functional connectivity in blindness by comparing resting-state fMRI data from sighted (n = 50) and early blind individuals (n = 33) using a data-driven hidden Markov model approach.

Early blind individuals exhibited increased fractional occupancy of two brain states characterized by bilateral somatomotor and left-lateralized occipito–frontal networks, whereas sighted participants more frequently occupied a state dominated by default mode and ventral attention networks. Connectivity states preferentially expressed by sighted participants showed reduced visual–somatomotor coupling in blind individuals, while states more frequently visited by early blind participants exhibited enhanced connectivity within the visual network and between occipital, fusiform, and prefrontal regions. Critically, the hyperconnectivity in blindness was largely explained by altered state occupancy, indicating that temporal dynamics, rather than connectivity patterns per se, drive reorganization.

Beyond temporal dynamics, visual deprivation altered the spatial organization of functional gradients derived from model-based, state-specific connectivity: occipital regions shifted closer to transmodal networks, while the overall low-dimensional geometry of latent states remained largely preserved. This suggests that early visual deprivation selectively reshapes the temporal expression and functional embedding of brain states without altering the core scaffold of large-scale dynamics.

Together, these findings demonstrate that blindness reshapes both the temporal and spatial structure of large-scale brain networks, highlighting the chronnectome as a critical substrate of experience-dependent plasticity.

**Significance Statement:** Blindness provides a unique opportunity to understand how the human brain reorganizes in the absence of typical sensory experience. Previous studies have relied on stationary functional connectivity, overlooking the role of temporal network dynamics. Using a data-driven Hidden Markov Model approach applied to resting-state fMRI, we show that early visual deprivation primarily alters the temporal dynamics of brain states. Crucially, the hyperconnectivity classically observed in early blind individuals emerges from changes in state occupancy rather than intrinsic alterations of connectivity patterns. Beyond temporal effects, we also demonstrate that visual deprivation reshapes functional gradients, with occipital regions shifting toward transmodal systems. Together, these findings reveal the chronnectome as a key mechanism of experience-dependent plasticity, while preserving a robust large-scale organizational scaffold.

## Introduction

The study of individuals who experience visual deprivation early in life provides a unique model to explore how the brain’s structural and functional architecture adapts following sensory deprivation. In early blind individuals (EBs), functional brain plasticity has been observed through either alteration of activity in non-deprived sensory regions, defined as intramodal plasticity (1–3) or enhanced processing of non-visual sensory information in deprived sensory areas, defined as crossmodal plasticity (4, 5) for review). The recruitment of occipital regions by nonvisual inputs in congenitally blind individuals underscores the experience-dependent nature of cortical organization. However, the functional specialization in occipital cortex in these individuals resembles that observed in sighted individuals and highlights the intrinsic constraints that shape and limit such plasticity (6–11).

To better understand how occipital regions in EBs are organized to support sensorimotor and cognitive functions, it is essential not only to examine their response properties (functional specialization) but also to explore how these regions are integrated into broader brain networks (functional integration) (12). A widely used approach for studying both short-and long-range functional brain interactions is functional connectivity (FC). FC quantifies the temporal correlation of functional activity between different brain regions, often expressed as pairwise Pearson’s correlation coefficients between time series (13). Traditional research on functional connectivity following early visual deprivation has consistently revealed increased or decreased connectivity involving occipital regions (see (14) for a review). Among the most reliable findings are a reduction in FC between occipital and tactile as well as auditory sensory cortical areas, concomitent to enhanced occipito-frontal connectivity in early blind people (15–22).

Traditional FC analysis assumes that interactions between brain regions remain stable over time. However, a more recent approach suggests that the wandering mind transitions between multiple functional modes, meaning that patterns of FC change over time despite the absence of a cognitive task (dynamic Functional Connectivity). Such fluctuations may provide insight into certain aspects of a neural system’s functional capacity (23, 24) and serve as a promising biomarker for disease (25, 26). From the perspective of dynamical systems theory, the brain can be understood as a system composed of various’attractors’, that are stable patterns of neural activity where brain states temporarily settle. Transitions between attractors allow flexible exploration of diverse network configurations (27). Computational models of whole brain network dynamics, based on tractography-derived realistic connectivity, suggest that resting-state activity operates at a critical bifurcation point on the edge of instability, exploring a variety of brain states (27–30). Minor changes in the system’s properties, such as shifts in the excitatory/inhibitory (E/I) balance, can lead to alterations in temporal brain dynamics (28, 31). Early sensory deprivation is known to trigger Hebbian and homeostatic plasticity mechanisms that may disrupt E/I balance (32, 33), leading to heightened occipital excitability and facilitating cross-modal reorganization (34). Consistent evidence from M/EEG and fMRI studies in early blind individuals indicates reduced occipital alpha, increased gamma activity, and elevated low-frequency fluctuations, reflecting altered inhibitory–excitatory dynamics (35–40). Building on this framework, we hypothesized that early visual deprivation systematically reshapes the temporal dynamics of large-scale brain states, providing a mechanistic motivation for investigating dynamic functional connectivity in blindness.

Dynamic functional connectivity (dFC) has most commonly been studied using the *sliding-window* approach, in which connectivity is estimated within consecutive segments of the fMRI time series (41, 42). This framework has been applied to clinical populations (20, 43–45), including early blindness (20), where increased temporal variability of occipito–temporal connectivity has been reported. This suggests a more flexible, multiple-demand role of the occipital cortex. However, window-based methods require arbitrary choices regarding window length and number, which can substantially influence results (26). The data-driven approach based on Hidden Markov Models (HMMs) overcome these limitations by identifying recurring brain states with distinct patterns of activity and functional connectivity, and by explicitly modeling their temporal evolution and transitions without imposing fixed windows (46–54). This method reveals the presence of a temporal hierarchy in which brain states are organized in two consistent and reproducible metastates: a metastate that covers higher cognitive functions and a metastate that covers sensorimotor systems and perception. The proportion of time spent visiting a specific brain state or metastate represents a subject-specific and heritable feature that can predict individual behavioral traits (55). Beyond these temporal metrics of the chronnectome, HMMs provide generative, state-specific connectivity estimates, offering a substrate for investigating how large-scale network dynamics relate to the spatial organization of cortical systems (56). Functional connectivity gradients have recently emerged as a principled framework to characterize the macroscale spatial organization of the cortex along continuous hierarchies from unimodal to transmodal regions (57, 58), and have been recently used to study experience-dependent plasticity in sensory deprivation (59, 60). However, it remains unknown whether and how visual deprivation affects the spatial organization of these cortical hierarchies at the level of dynamic brain states.

In this study, we assessed dFC in resting-state fMRI data from early blind and sighted individuals to address two main objectives. First, we examined whether early visual deprivation leads to systematic alterations in the temporal dynamics of functional connectivity. Second, we asked whether such dynamic changes can help explain previously reported differences in stationary functional connectomes, in particular the unexpected reduction in connectivity between occipital and sensorimotor regions (16, 17). To this end, we applied HMMs to characterize group differences in chronnectome patterns and to relate these temporal features to large-scale topographical properties of the functional connectome.

Building on this framework, we further exploit the generative nature of the HMM to move beyond temporal descriptors of network dynamics. Specifically, we tested two complementary hypotheses: (i) visual deprivation alters the organization of functional gradients when they are estimated from model-based, state-specific connectivity patterns rather than stationary FC; and (ii) visual deprivation reshapes the low-dimensional geometry of the latent states underlying dynamic brain activity. By jointly characterizing spatial hierarchy and state-space geometry, our approach links experience-dependent plasticity to both the temporal and spatial structure of large-scale brain dynamics.

## Results

### State characterization

To delineate the spatial layout of each hidden state, we derived a mean activation vector for each state. These vectors, spanning across 100 parcels, encapsulate the activation levels of each parcel when a specific state (K) is active. In Fig. 1A, we visually depict the spatial arrangement of this vector, assigning the same value to all vertices within each parcel. Notably, the data used to infer the HMM were standardized to ensure a zero mean and unit standard deviation across time for each parcel. Consequently, an activation level of zero corresponds to the mean activation of the respective parcel. Negative activations (depicted in cool colors) indicate below average BOLD activity, while positive activations (shown in warm colors) denote above average BOLD activity.

**Figure 1.**
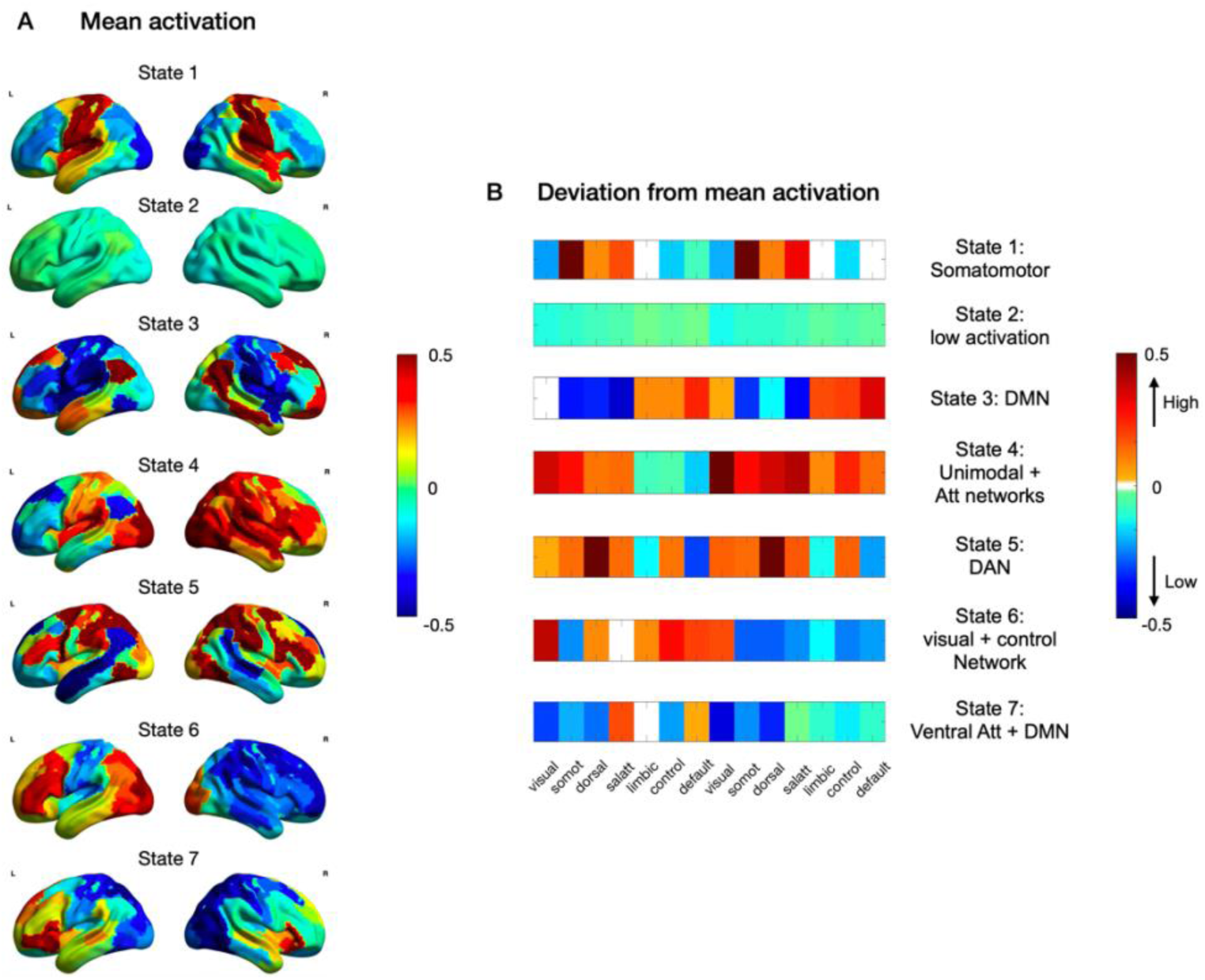
States characterization. **A.** Whole brain maps representing the mean activation on the lateral brain surface. Each parcel is color-coded according to the activation extracted when the specific state is active. **B.** Deviation from the mean level of activation for each resting state network and hemisphere (RSN order: visual, somatomotor, dorsal attentional, salience, limbic, control and default mode network). Warm colors indicate activations higher than the mean; cold colors indicate activations lower than the mean.

To clarify which resting state network (RSN) most defines each hidden state, we computed a summary measure for each hemisphere and RSN. This measure quantifies the extent of deviation of each RSN from the average level of activation. In our visualization (Fig. 1B), warmer colors represent higher positive deviations from the mean, while cooler colors represent higher negative deviations from the mean. Among all brain states, states 2 and 4 stand as the primary extremes: the former exhibits generally low mean activation, while the latter displays high activation primarily within unimodal and attentional networks. Additionally, state 1 showcases elevated activation mainly in regions belonging to the somatomotor network, while state 6 manifests heightened activation in visual and frontal regions. The remaining states predominantly feature activation in multimodal regions, including the default mode network (DMN) in state 3, the dorsal attentional network in state 5, and a combination of the ventral attentional network and DMN in state 7.

### Altered brain state dynamics in blindness

We conducted between-group comparisons on three pivotal metrics defining the *chronnectome*: 1) Fractional Occupancy (FO), indicating the proportion of time spent visiting a state; 2) Mean Lifetime, denoting the number of timepoints per visit; and 3) Switching Rate (SR), representing the frequency of transitions between states. In Fig. 2, we present the results for all metrics.

**Figure 2.**
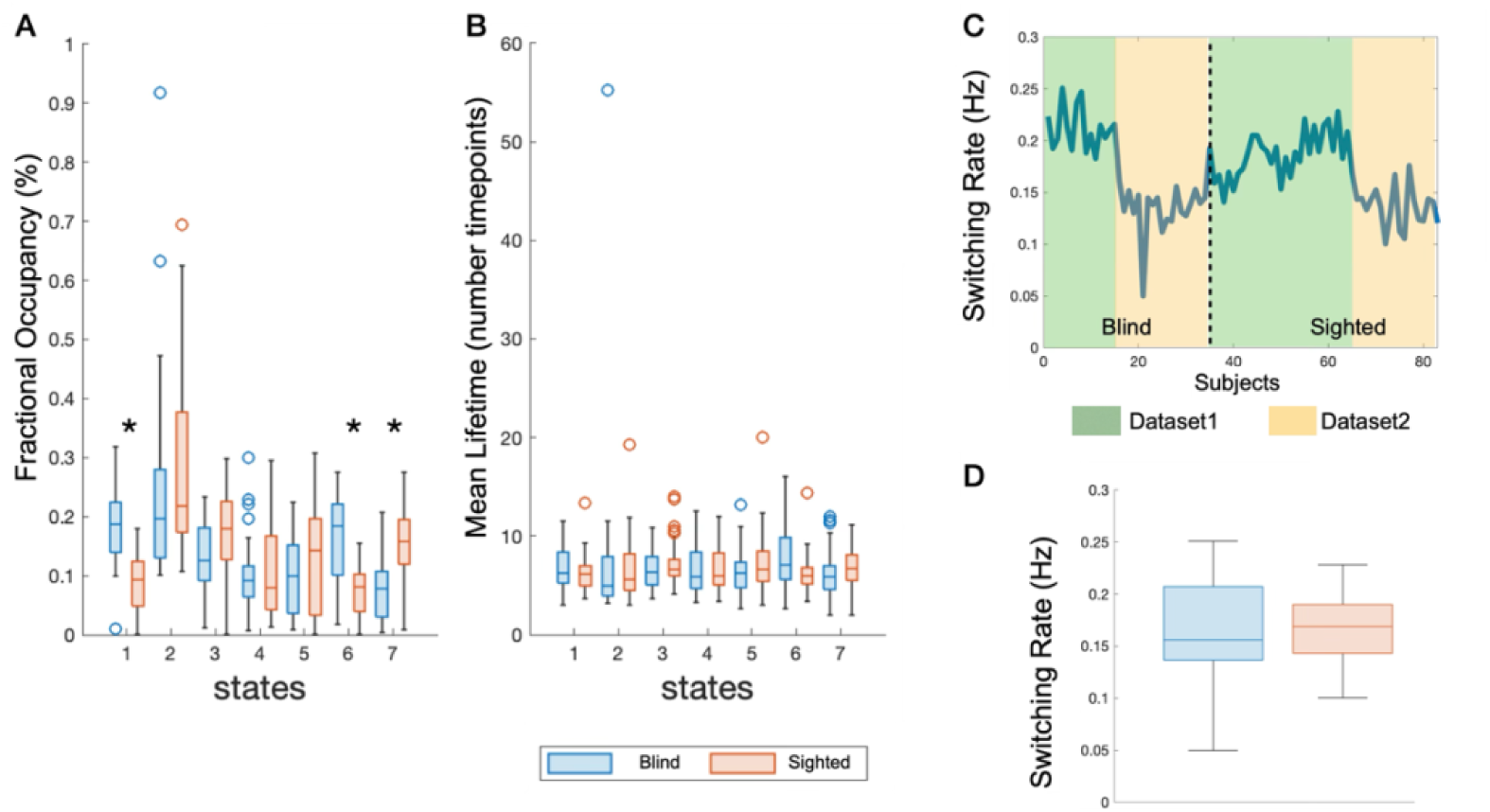
Differences in the *chronnectome* between blind and sighted individuals. **A.** Fractional Occupancy for each group, as a function of state. **B.** Mean Lifetime for each group, as a function of state. **C.** Switching rate (Hz) for each subject (upper), with green background indicating subjects belonging to the dataset from Trento and orange background indicating subjects belonging to the dataset from Montreal, and for each group (bottom). Significant results after FDR correction are highlighted with *.

Significant disparities emerged in Fractional Occupancy (Fig. 2A), with blind people showing a propensity to visit states 1 and 6, whereas controls exhibited a propensity to visit state 7 (FDR-p=0.004). It is crucial to contextualize these findings within the characterization of the states to glean deeper insights. States 1 and 6 primarily exhibit activation in unimodal regions, with state 1 showing heightened activity in somatomotor regions and state 6 in visual areas (see Fig. 1). Furthermore, state 6 entails activation of regions within the Default Mode and Control networks, predominantly lateralized to the left hemisphere. In contrast, controls tend to favor state 7, characterized by activation in regions of the Default Mode and Ventral Attentional networks, devoid of involvement from unimodal regions. Upon conducting separate analyses on the two datasets, we observed similar results (see Fig. S4), confirming that these primary findings exhibit robustness across datasets. In addition, we observed a discrepancy in state 3 (FDR-p = 0.009), wherein controls displayed higher Fractional Occupancy when analyzing the dataset from Trento. Interestingly, in 2 out of 3 states in which we did not find any significant between group difference (state 4 FDR-p = 0.51 - state 5 FDR-p = 0.16), we observe a clear activation of bilateral occipital areas, together with unimodal and multimodal regions.

No significant disparities were detected in the mean lifetime metric (Fig. 2B), suggesting that both groups allocated a similar number of timepoints per visit, irrespective of differences in Fractional Occupancy. This implies that even in states exhibiting higher Fractional Occupancy in either blind or sighted individuals, the average duration of each visit remained consistent across groups. Upon conducting separate analyses on the two datasets, we observed congruent results (see Fig. S4), with the sole exception of a significant difference in state 3 (p_FDR_ = 0.006) when analyzing the dataset from Trento, indicating a longer visit duration in sighted compared to blind individuals.

We did not find any difference in terms of switching rate (FDR-p=0.115), with blind and sighted individuals switching among states every 0.169Hz (5.91 seconds) and 0.167Hz (5.98 seconds) respectively (Fig. 2D). The examination of the switching rate trend across subjects (Fig. 2C) revealed generally higher values in the first than in the second dataset, as well as a bigger between-group difference when considering only the first dataset. Indeed, upon conducting separate analyses on the two datasets, we observed a significant difference only when considering the dataset from Trento, with significantly higher switching rate in blind (0.209Hz) than sighted individuals (0.185Hz - FDR-p = 0.027). Thus, any potential between-group differences in this regard may be attributed more to sample characteristics or scanner type rather than early visual deprivation.

In our analysis, we explored various model orders ranging from 5 to 9 states, as detailed previously (Fig. S5). Interestingly, consistent findings emerged when evaluating between-group disparities in Fractional Occupancy (FO). The most consistent result across models was an increased FO in blind individuals within a state characterized by bilateral activation of somatomotor and auditory regions. Moreover, blind people exhibited elevated FO in states featuring robust activation of the visual network, often in conjunction with fronto-parietal regions. Conversely, controls tended to display higher FO in states characterized by multimodal region activation and cortical hubs, notably regions associated with the Default Mode Network. These findings highlight that our conclusions are not purely driven by the order chosen.

### Transition Probabilities

An additional feature of the Chronnectome is represented by the matrix of transition probability (K x K matrices, where K is the number of states). We extracted these matrices at the subject level, excluding probabilities of persistence (diagonal of the matrix), by using the function *getTransProbs* of the HMM-MAR toolbox, and computed the median values separately for blind and sighted individuals (see Fig. 3A). Fig. 3B illustrates the hierarchical clustering (using Ward’s algorithm) on the resulting transition probability matrices, for blind (top) and sighted (bottom) individuals. Remarkably, both groups showed similar patterns, with an increased likelihood of transitioning to state 2 from any other state. This state is characterized by overall low brain activation, close to the mean, and replicates previous findings suggesting that such a low-activation state may serve as a transitional hub in resting-state fMRI (61, 62). In addition to this overarching trend, when considering the top 20% highest probabilities in both groups, notable distinctions emerge (see Fig. 3C). Blind people displayed a greater likelihood of transitioning from state 7 or 2 to state 1, whereas controls exhibited a higher probability of transitioning from state 2 to either state 3 or state 7. These findings align well with the previously observed patterns of higher FO in blind subjects in state 1 and higher FO in sighted in states 3 and 7.

**Figure 3.**
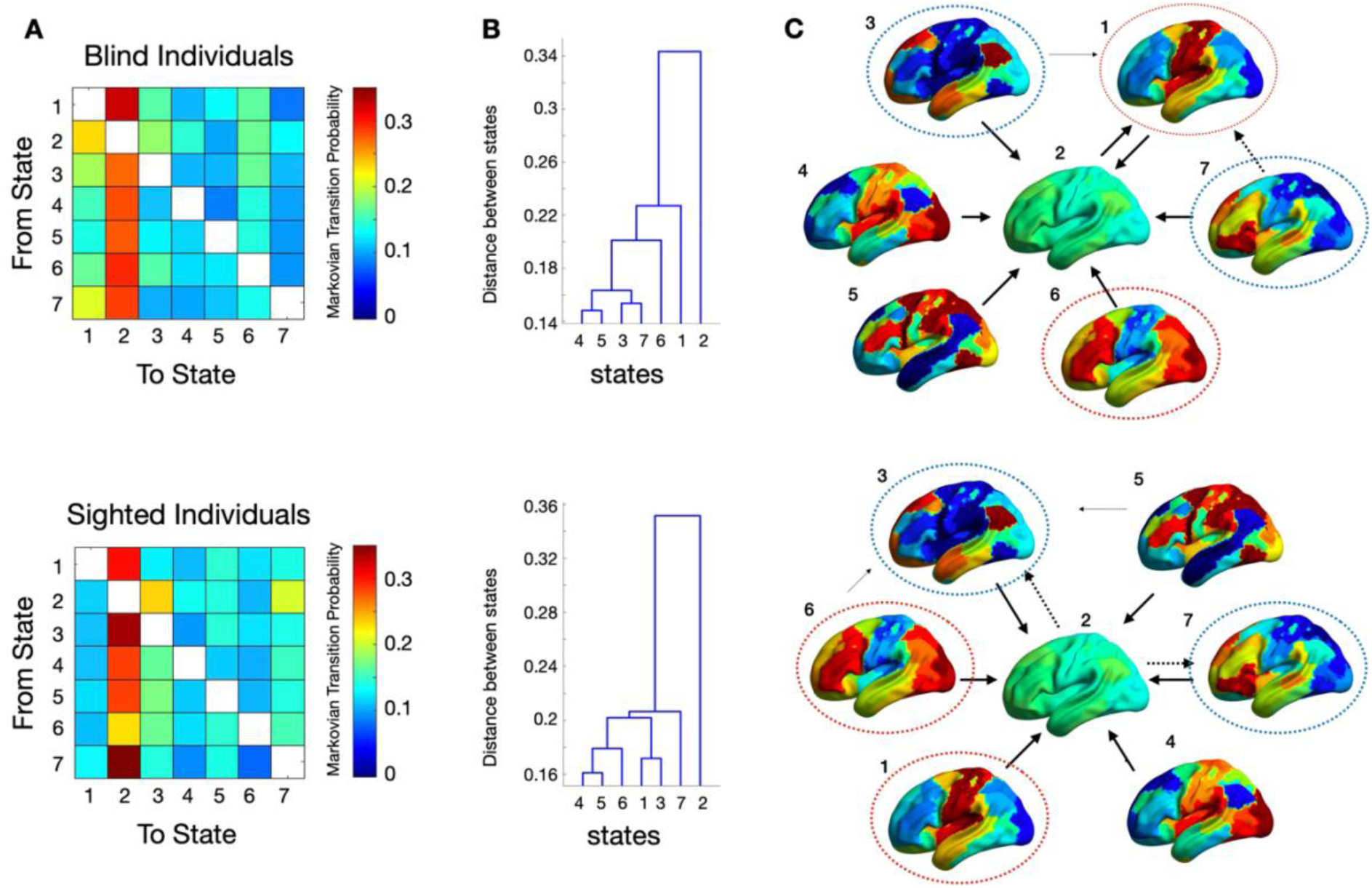
Transition probability matrices in blind and sighted individuals. **A.** Median transition probability matrices for blind (top) and sighted (bottom) individuals. The color scale indicates the Markovian transition probability from-to a specific state. **B.** Dendrogram showing the clusters of states based on the median transition probabilities. **C.** Graph showing the 20% higher probabilities between states.

### Functional connectivity

When referring to the stationary functional connectome, we considered two subject-level FC representations: (1) an empirical FC matrix, estimated as the full correlation between pairs of parcels, and (2) a model-based time-averaged connectome, computed as the weighted sum of the FC patterns of all HMM states.

We first extracted the empirical stationary FC by calculating the full correlation between all parcel pairs. This analysis enabled us to verify that our data reproduced the well-established group differences reported in the literature. Specifically, we observed two main patterns: enhanced connectivity between occipital and frontal regions in EBs, and increased connectivity between occipital and somatomotor regions in SCs, confirming previous findings (see Fig. 4A) (17, 40, 63–65). Establishing this expected stationary FC pattern in our dataset provided a critical baseline for the subsequent analyses. Interestingly, the model-based stationary functional connectivity (rs-FC) showed a highly similar pattern. Blind individuals exhibited reduced rs-FC between visual and somatomotor regions and increased rs-FC between visual and prefrontal regions. In addition, we observed stronger rs-FC in blind participants between visual areas and both the bilateral precuneus (a core region of the DMN) and the fusiform gyrus (see Fig. 4B).

**Figure 4.**
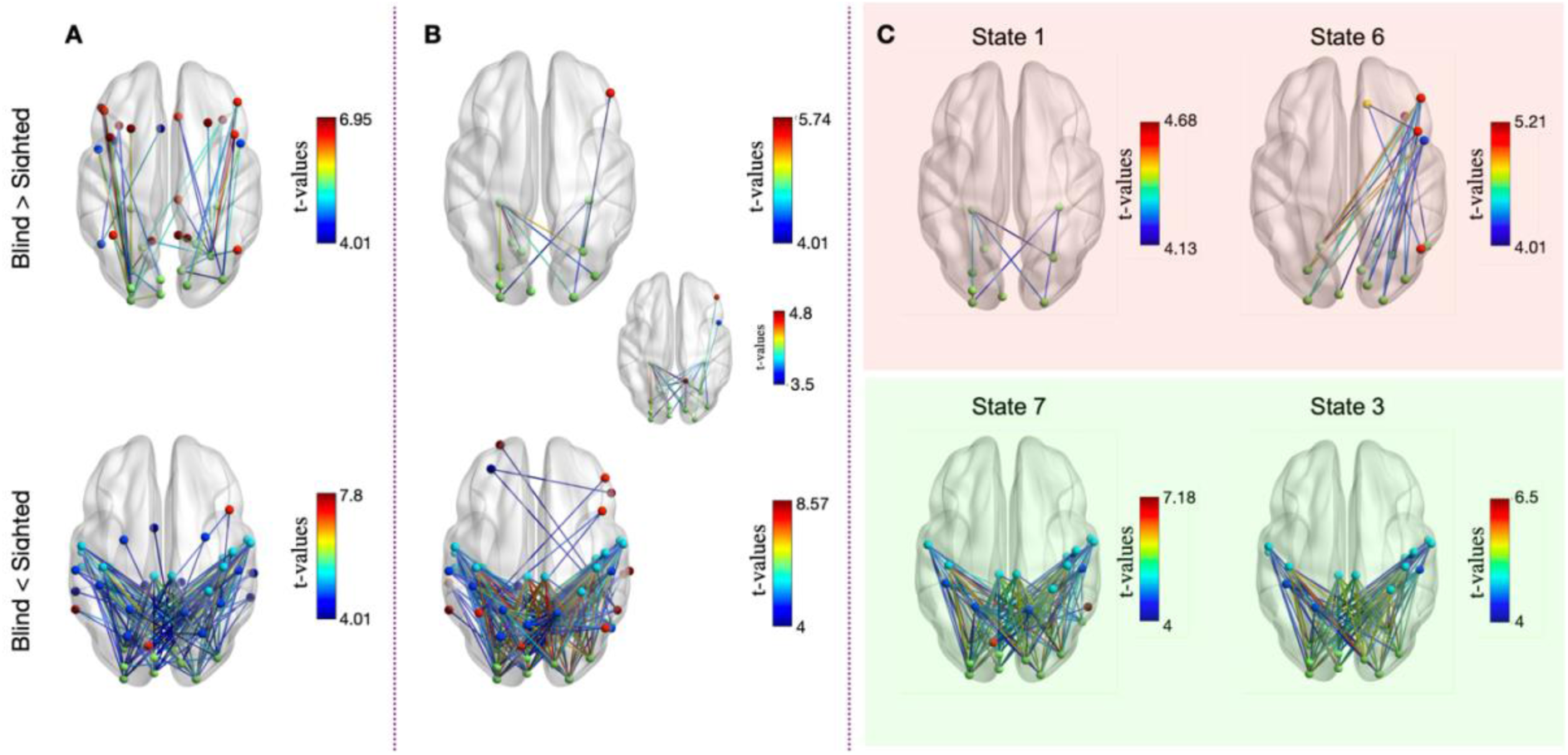
Network-level between-group analysis. Between-group differences in the functional connectome for the (A) empirical stationary FC, (B) time-averaged functional connectome and (C) the time-varying FC. In each figure, we show the connections that are significantly different between groups, color-coded based on their t-values. The red background indicates states with FO higher in blind individuals; the green background indicates states with FO higher in controls.

When examining the time-varying functional connectome (Fig. 4C), we identified distinct connectivity patterns across dynamic states. In four out of seven states, blind participants showed reduced functional connectivity between visual and somatomotor regions compared to controls. Reductions in connectivity were most evident in states with higher FO in controls, specifically states 3 and 7, and in state 2 (Fig. S6), which showed globally low activation and moderately increased connectivity within the Default Mode and Control Networks. In contrast, three states exhibited higher functional connectivity (FC) in blind participants. States 1 and 5 showed increased FC within the visual network, particularly involving the Vis1 parcel of the Schaefer Atlas (Fig. 4B, S6), likely overlapping with the posterior parahippocampal and fusiform gyri as defined by the Harvard–Oxford Cortical Atlas. State 6 was characterized by enhanced FC between occipital and right prefrontal regions, reflecting stronger inter-network coupling. Notably, states 1 and 6 were more frequently visited by blind individuals. Importantly, states 1, 5, and 6 - those associated with stronger rs-FC in the blind - corresponded to the increases observed in the stationary FC analysis. In contrast, states 2, 3, and 7 - those with higher FO in controls - mirrored the stronger rs-FC observed in controls in the stationary analysis. Together, these findings indicate that group differences identified by stationary FC analysis likely reflect not only changes in rs-FC strength but also differences in the temporal dynamics, namely the amount of time spent in specific connectivity states.

To quantify the contribution of each latent state to stationary FC, we performed an edge-wise regression analysis in which the dependent variable was either the model-based or the empirical FC and the regressors of interest were the HMM-states FC weighted for the FO. For the model-based FC, the model explained approximately 99.1% of the variance in stationary FC, which exceeded chance levels for 99.75% of all edges. The median edge-wise correlation between the predicted and observed FC was 0.99 (Fig. S7A). This high correspondence is expected, given that the model-based stationary FC is computed as the weighted average of the state-specific FC patterns. For the empirical stationary FC, the HMM-based model captured a substantial portion of the variance (adjusted R2 = 33.2%) and predicted 93.8% of edges significantly above chance, with a median edge-wise correlation of 0.64 (Fig. S7B). Interestingly, in both cases the variance explained decreased when we did not weight the FC of each HMM state for its temporal dynamic (FO, adjusted R2 empirical FC = 91.4%; adjusted R2 model-based FC = 31.7%). This indicates that dynamic HMM-derived FC provides a reasonably good approximation of the stationary FC structure and that FO adds significant information to the estimation.

### Two prevalent brain states account for stationary FC differences in early blindness

To assess the contribution of each state to the between-group differences observed in time-averaged FC, we computed state-specific, FO–weighted FC matrices for each subject. For each state, FO-weighted FC matrices were summarized at the group level using the median across subjects, separately for EB and SC. We then computed a between-group difference matrix (Δ = EB − SC). Each state was summarized by (i) the mean Δ across all off-diagonal edges and (ii) the fraction of edges showing higher FC in EB (Δ > 0) or in SC (Δ < 0). We observed a positive Δ in states 1, 4, and 6, with more than 70% of edges showing higher FC in EBs (95.4% in state 1, 71.2% in state 4, and 87.9% in state 6). In contrast, all other states showed a negative Δ, indicating higher FC in controls, with states 2, 3 and 7 showing the highest percentage of edges favoring controls (87.7% in state 2, 73.7% in state 3 and 87.6% in state 7).

Using a leave-one-state-out approach to reconstruct the time-averaged FC, we found that the between-group differences reflecting hyperconnectivity in EB were primarily driven by states 1 and 6 (Fig. 5, upper row). Removing either of these states completely removed the pattern of hyperconnectivity observed in EBs. In contrast, removing any other state had little effect on the significance of most edges. Moreover, when removing either state 2 or 7, additional connections showing EB hyperconnectivity emerged, mainly between occipital and frontal regions, similar to edges observed in the empirical FC (Fig. 4A). These states were more frequently visited by controls, with a significant difference in FO observed only for state 7.

**Fig. 5.**
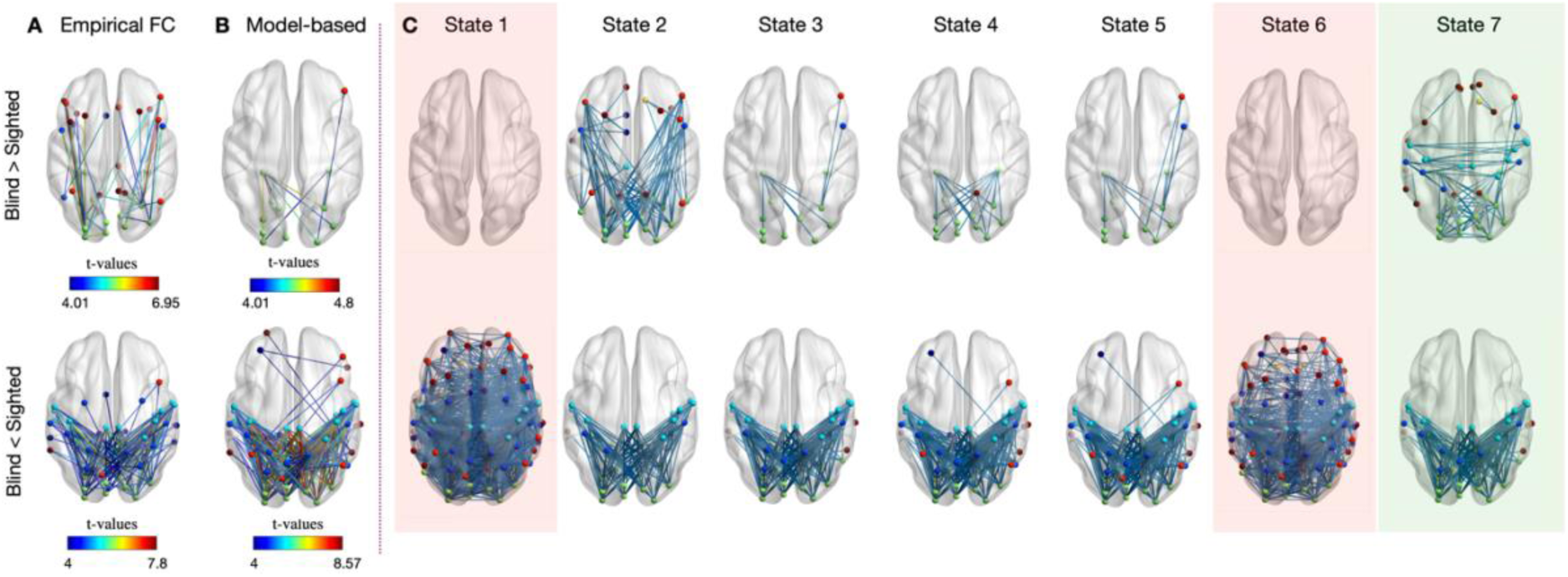
Patterns of functional connectivity in the (A) empirical FC, (B) time-averaged FC and (C) reconstructed time-averaged FC using a leave-one-state-out approach. Results are shown for both contrasts: Blind > Sighted (upper row) and Sighted > Blind (bottom row). In the empirical and model-based, time-averaged FC (A and B), edges are color-coded according to t-values, whereas in the reconstructed time-averaged FC (C) all edges are shown in blue to indicate which connections are maintained across reconstructions. Red and green backgrounds indicate states in which FO is significantly higher in EBs (red) or in controls (green).

Considering hyperconnectivity in SCs, the main between-group differences were primarily driven by states 2, 3, and 7. However, removing any of these states (Fig. 5, bottom row) led only to a partial decrease in the number of edges showing increased connectivity in SC, with 50.2% (state 2), 32.9% (state 3), and 33.7% (state 7) of edges losing significance. In contrast, removing EB-dominant states 1 and 6 had little effect on most edges but revealed several additional significant connections (73.6% for state 1, 58.6% for state 6), mainly involving frontal regions. The emergence of these additional hyperconnected edges following the exclusion of specific states should be interpreted as a consequence of reweighting the remaining state contributions to the reconstructed time-averaged FC. Certain states can reduce the detectability of between-group differences by emphasizing alternative or weaker connectivity patterns; removing them can unmask latent group differences that are otherwise averaged out in the full time-averaged representation.

Taken together, these findings confirm that states 1 and 6 primarily drive EB hyperconnectivity, whereas multiple states contribute to the pattern of hyperconnectivity in SCs.

### Temporal dynamics underlies functional connectivity alterations in early blindness

The main goal of this analysis was to disentangle the contributions of the temporal dynamics of brain states, quantified by fractional occupancy (FO), from the contribution of state-specific functional connectivity (FC) to the between-group differences observed in stationary FC. We focused on states 1 and 6, which primarily drove EB hyperconnectivity, and state 7, which primarily drove SC hyperconnectivity.

For states 1 and 6, EB hyperconnectivity was largely determined by the temporal component (Fig. 6). When FO values of controls were assigned to EBs, the hyperconnectivity disappeared in all permutations, even after reducing the threshold to 3.5. The same result was observed when using the mean FO of controls. In contrast, when the functional connectivity (FC) patterns of controls were reassigned to EBs while keeping the original FO values, between-group differences persisted in ∼10% of permutations (up to ∼45% for State 1 and ∼20% for State 6 when using a lower threshold of 3.5), mainly involving occipital regions in the left hemisphere. Similarly, assigning the mean FC of controls to EBs did not eliminate the hyperconnectivity, indicating that differences in intrinsic FC patterns did not drive the effect. Together, these results show that hyperconnectivity in states 1 and 6 in EBs is primarily driven by differences in temporal expression (FO), rather than by the spatial pattern of connectivity.

**Figure 6.**
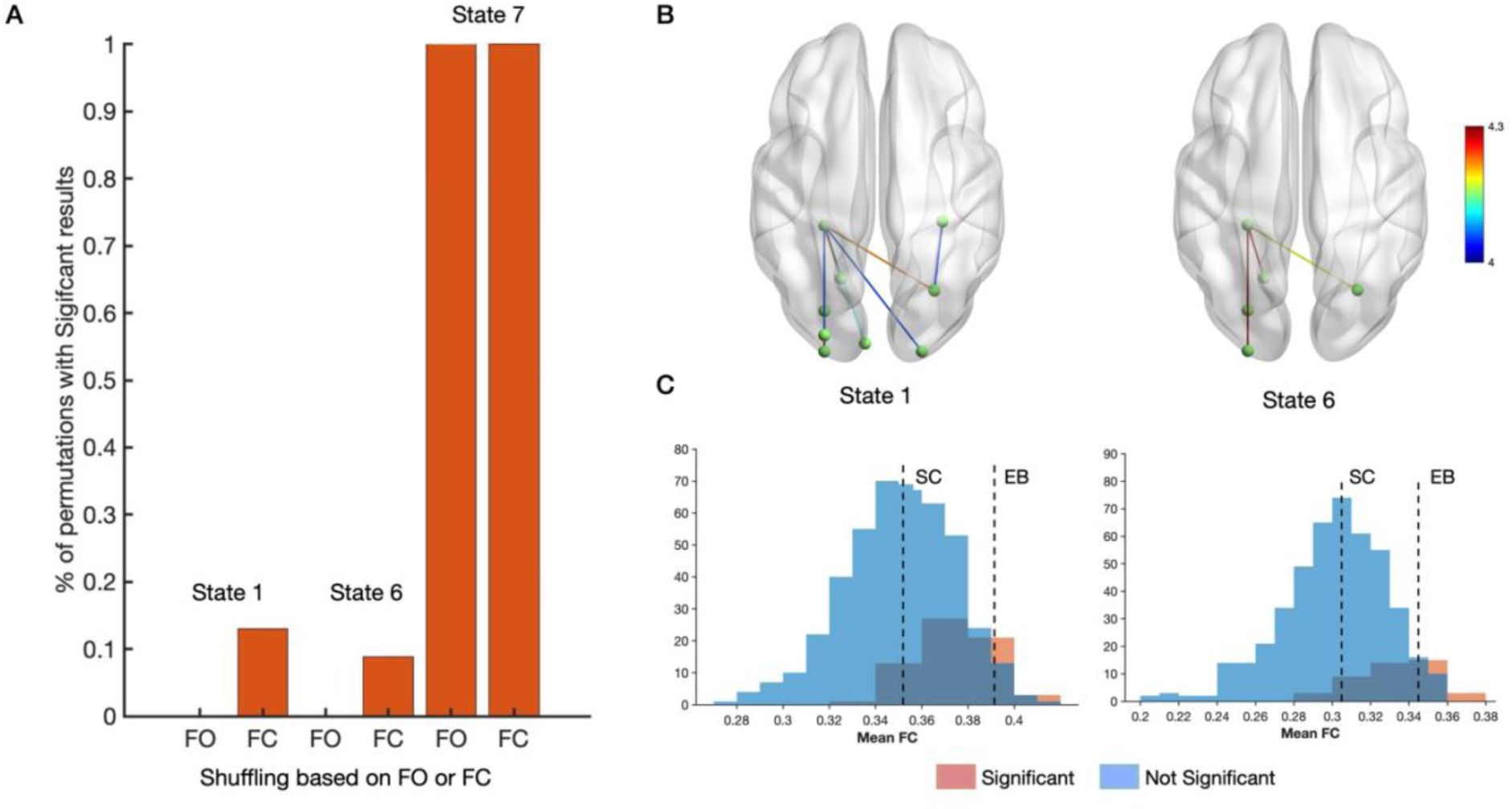
Temporal and spatial contributions of HMM states to hyperconnectivity in EB participants. **A.** Percentage of permutations showing significant hyperconnectivity in EBs for State 1 and 6, and in SCs in State 7, after shuffling either fractional occupancy (FO) or functional connectivity (FC). **B.** Pattern of significant hyperconnectivity in EBs, shown for the permutation with the largest number of significant edges. **C.** Distribution of the mean FC in EBs for permutations with (red) or without (blue) significant between-group differences. Dashed lines indicate the empirical mean FC for each group.

When considering state 7, one of the states contributing the most to the pattern of hyperconnectivity in SCs, both FO and FC manipulations did not reduce the between group differences. This result confirms our previous findings (*State-wise contributions to stationary functional connectivity),* showing that many connections remained significant even after removing state 7, indicating that SC hyperconnectivity is mediated by multiple states rather than a single state. These findings suggest that the between-group differences in SC are more distributed across the temporal dynamics and intrinsic connectivity of several HMM states.

### Occipital cortex shifts toward transmodal networks

In a seminal study, Margulies et al. (2016) applied a nonlinear dimensionality reduction algorithm to the resting-state functional connectomes of 820 individuals, revealing the principal axes of variance in brain connectivity. They showed that the first gradient dissociates unimodal from transmodal regions (e.g., the default mode network, DMN), whereas the second gradient differentiates visual from somatosensory systems. These gradients, described as the ‘intrinsic coordinate system’ of the human brain (58), have been linked to variations in brain structure (66–68), gene expression (69), and information processing (58). Here, we assessed the effect of early sensory deprivation on (1) the spatial organization of functional gradients, mainly focusing on the distance between the visual network and other resting-state networks, and (2) the alignment of Hidden Markov Model (HMM) states along the two main gradients. We focused on the first two gradients, which together captured at least 60% of the total embedding contribution in both groups (Fig. 7A).

**Fig. 7.**
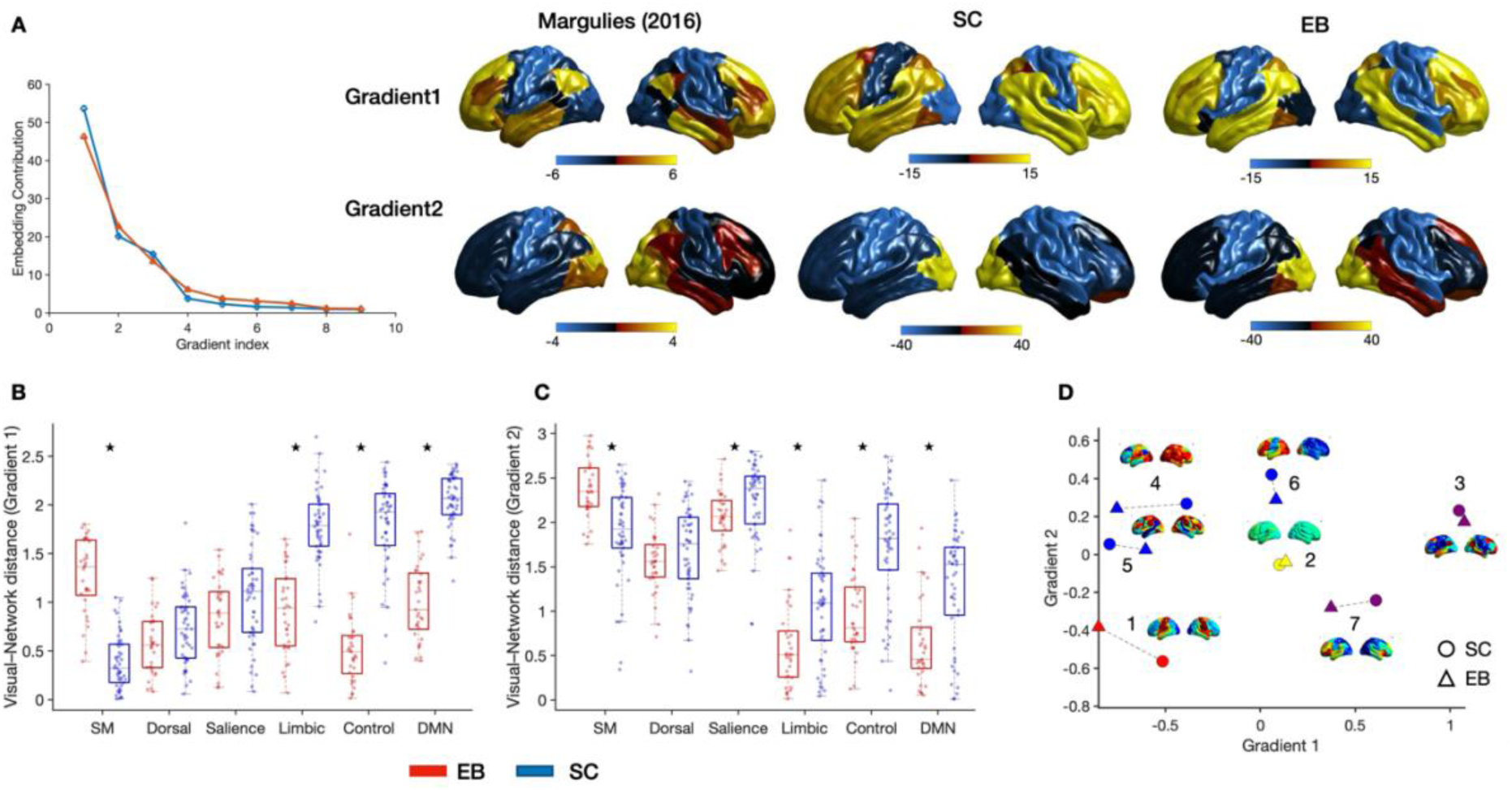
Functional gradients and HMM state projections. **A.** Embedding Contribution as a function of the Gradient Index. Spatial maps of cortical gradients 1 and 2 projected onto the lateral brain surface for three datasets: reference data from Margulies et al. (2016), sighted controls (SC), and early blind individuals (EB). **B–C.** After projecting each subject-specific model-based FC into the group-specific functional gradient, we calculated the distance between the visual network and all other resting-state networks (RSNs) for EBs (red) and SCs (blue). Distances are shown separately for Gradient 1 (B) and Gradient 2 (C). Asterisks indicate the significant results after correcting for multiple comparisons (FDR). **D.** Projection of the mean activation of Hidden Markov Model (HMM) states onto the low-dimensional manifold defined by the reference cortical gradients. Each state is represented for EBs (triangle) and SCs (circle), and color-coded according to the associated activation map: red for state 1, predominantly activating sensorimotor regions (SM); blue for states involving occipital regions; yellow for the low-activation state; and purple for states not activating any sensory regions. The group mean activation map is provided for each HMM state.

In general, we observed a high correlation between the reference and the population specific gradients (r = 0.68 Gradient 1, r = 0.9 for Gradient 2 in SCs; r = 0.85 Gradient 1, r = 0.91 for Gradient 2 in EBs). When plotting the cortical gradients derived from the model-based stationary FC for SCs and EBs, we observed clear differences along the unimodal–transmodal gradient (Gradient 1; Fig. 7A). Specifically, occipital regions in EBs were shifted toward transmodal areas, lying between the DMN and sensorimotor regions in gradient space. After projecting each subject’s model-based stationary FC onto the group-level gradients, we observed several between-group differences. Along the unimodal-transmodal gradient (G1), we did not observe a significant difference in the gradient range, indicating a similar degree of functional heterogeneity between groups (p = 0.53, r =-0.07). However, we found a significant reduction in the distance between the visual network and the default mode network (DMN) in EBs compared to SC (p < 0.001), indicating that occipital regions were positioned closer to transmodal regions along the principal gradient (Gradient 1; Fig. 7B). More generally, these results indicate that sensory deprivation is associated with a reorganization of the unimodal–transmodal gradient, with occipital regions moving further away from sensorimotor regions and closer to limbic, control, and default mode networks. These group differences were associated with large effect sizes, as quantified by the rank-biserial correlation (|r| range = from 0.74 to 0.82), with particularly strong effects for distances between visual and SM (r = 0.77), limbic (r =-0.74), control (r =-0.79), and DMN (r = - 0.82) networks. Distances between the visual network and attentional networks (dorsal and salience) were not significant (r =-0.22). Along the visual–sensorimotor gradient (Gradient 2), we observed a significant reduction of the gradient range, possibly indicating functional homogenization (p = 0.0001, r =-0.44) following sensory deprivation. Moreover, EBs exhibited a significant increased distance between visual and sensorimotor networks (p < 0.001, r = 0.48), while distances between visual and higher-order cognitive networks (salience, limbic, control, DMN) were generally reduced, except for the dorsal attentional network (Gradient 1; Fig. 7C). In this gradient, group differences showed moderate effect sizes (ranging from r = 0.31 for salience to r =-0.54 for control network). Taken together, these results indicate a shift of occipital regions toward transmodal networks along both the two cortical gradients.

Considering each latent state separately revealed distinct group differences along both principal gradients (Fig. S8). Along Gradient 1, we found that numerous significant between-group differences were observed across states. Of particular relevance, the strongest reduction in distance between occipital and transmodal regions in EBs was seen in states 1 and 6 (DMN: r =-0.83,-0.84; Control: r =-0.83,-0.84), whereas state 7 showed a reversed pattern, with EBs exhibiting increased distance to transmodal regions (DMN: r = 0.3). For Gradient 2, we observed that the main effect observed was consistent across all states, with EBs showing greater separation between the visual and somatomotor networks relative to controls. The largest effect was observed in state 7 (r = 0.73), indicating a pronounced increase in visual–somatomotor distance in EBs. While multiple state-specific effects reached significance, these highlighted results capture the most relevant reorganization of functional gradients associated with visual deprivation.

### Brain state dynamics map onto low-dimensional functional gradients

We projected HMM-derived latent states onto the principal cortical gradients to examine whether early visual deprivation reorganizes functional brain dynamics in low-dimensional space. Across both groups, Gradient 1 largely dissociated states dominated by sensorimotor versus transmodal (DMN) activation, whereas Gradient 2 primarily separated states characterized by sensorimotor versus occipital activation (states 4–6; Fig. 7D). Low-activation states (e.g., state 2) were positioned near the center of the manifold. These observations are consistent with previous work (Song et al., 2023), showing that HMM states align along principal gradients, with Gradient 1 reflecting unimodal–transmodal dissociation and Gradient 2 reflecting somatomotor–visual dissociation, indicating that temporal dynamics of functional connectivity are constrained by cortical gradient architecture. Notably, early visual deprivation was associated with systematic shifts in specific states. The Euclidean distances between EB and SC state coordinates ranged from 0.036 to 0.383 across states, with the largest divergences observed in states 1 (0.383) and 4 (0.369), indicating that these states are most reorganized in EBs relative to SCs. In contrast, state 2 exhibited minimal difference (0.036), consistent with its low-activation profile near the manifold center. Overall, these results suggest that early sensory deprivation selectively reshapes the geometry of latent neural states, particularly for states involving deprived and not-deprived sensory networks, while preserving the general gradient-based organization of functional dynamics.

## Discussion

Early blindness alters the structural and functional architecture of the brain. Previous studies have documented changes in the functional connectome (17, 63, 64, 70). In this study we assessed whether early blindness could lead to changes in the temporal dynamics of functional connectivity (FC), by applying Hidden Markov Models to resting state fMRI data of early blind (EBs) sighted (SC) individuals.

The first goal was to assess whether early visual deprivation could lead to alterations in the temporal dynamics of the functional connectome. We observed significant differences between EBs and SCs in the frequency of occupying specific brain states (Fractional Occupancy). Interestingly, the overall switching rate between states was similar across groups, suggesting that early sensory deprivation does not alter the multi-stability feature of the system. **Blind individuals** tended to visit more frequently two brain states characterized by increased cross-talk among auditory and somatomotor regions and between occipital and frontal cortices. The elevated occurrence of states engaging unimodal sensory and frontal systems may reflect baseline fluctuations that predispose both deprived and non-deprived sensory regions to heightened recruitment during perceptual and cognitive tasks following visual deprivation (71–78). Interestingly, the increased engagement of unimodal sensory regions may reflect the enhanced processing of exteroceptive (e.g., auditory and tactile) or interoceptive (e.g., heartbeat) information previously reported following early visual deprivation (79–83). Within the state dominated by bilateral somatomotor activity, EBs exhibited increased connectivity within the visual network, primarily among occipital regions and ventral temporal areas, including the parahippocampal and fusiform gyri, which are implicated in spatial context processing and high-level object recognition. In the occipito–frontal state, we observed enhanced coupling between early visual and prefrontal cortices, consistent with prior reports of strengthened interactions between deprived visual cortex and higher-order cognitive systems supporting language, executive control, and working memory (22, 73, 84). Taken together, these findings support the hypothesis that portions of the occipital cortex acquire properties of a multiple-demand system (85), with stronger coupling to frontal regions reflecting their integration into domain-general cognitive processing (20). **Sighted individuals** more frequently visited a state characterized by the activation of regions belonging to the Default Mode and Ventral Attentional networks. One of the main regions activated in this state was the medial prefrontal cortex, a core hub of the default mode network (86) implicated in mental imagery (87–90). Interestingly, in this state, we observed a higher visual–auditory/somatomotor connectivity in controls, consistent with previous stationary FC findings (17, 63–65). We therefore speculate that this state may reflect imagery-related coactivation and enhanced coupling among sensory areas in sighted individuals, and that its reduced occurrence in blindness indicates diminished reliance on visual imagery and reorganization of sensory–motor interactions. This pattern suggests that occipital connectivity may be task-dependent: in SCs the external sensory input may suppress occipital activity during tasks (91), reducing cross-modal coupling (92), whereas in EBs this pattern could be reversed, leading to reduced coupling at rest and increased connectivity during sensory processing (20).

The second goal was to test whether the alterations in functional connectivity traditionally reported following sensory deprivation are driven primarily by changes in the temporal dynamics of brain states, rather than by the connectivity patterns themselves. Crucially, our results clearly show that the hyperconnectivity observed in EBs is driven entirely by the two states they occupy most frequently, and that fractional occupancy is its primary determinant. This indicates that group differences in stationary resting-state FC mainly reflect how often specific connectivity configurations are expressed, rather than differences in the configurations themselves. By contrast, hyperconnectivity in SCs emerges across multiple states and reflects a combination of temporal and spatial features. These findings fundamentally shift the interpretation of connectivity alterations in blindness, moving from a purely spatial reorganization of network architecture to a dynamical reweighting of the brain’s functional repertoire, in which altered time allocation across latent states plays a central role. This temporal perspective provides a mechanistic explanation for previously reported stationary FC effects and underscores the need to move beyond static connectivity maps toward models that explicitly capture the brain’s evolving functional organization. Importantly, recognizing the central role of temporal dynamics may have direct translational implications. State prevalence may serve as an individualized biomarker of functional organization and plasticity, while interventions based on neurofeedback (93) or stimulation could be designed to selectively promote or suppress specific brain states, thereby modulating large-scale network function in a targeted and mechanistically informed manner.

In the last part of this study, we examined how alterations in large-scale network dynamics relate to the spatial organization of functional gradients. To this end, we projected the model-based functional connectivity estimated by the HMM onto the principal gradients (G1 and G2) and tested for group differences in functional differentiation (gradient range) and in the segregation of the visual network. Unlike previous work based on conventional stationary FC, this approach isolates connectivity patterns that are explicitly grounded in the brain’s latent dynamic states. We found that early visual deprivation does not disrupt the global hierarchical organization of the principal gradients. However, network-level analyses revealed a selective reconfiguration: occipital regions were positioned farther from other sensory networks and closer to transmodal regions, primarily belonging to the Control and Default Mode Networks. This shift indicates that the functional connectivity profile of occipital cortex in blindness becomes more similar to that of transmodal rather than unimodal regions. These findings support the idea that genetically guided structural constraints establish the cortical hierarchy, while sensory experience reshapes its functional expression through adaptive, fine-grained reorganization (8, 94, 95). Crucially, this reorganization was not uniform across time. By repeating the gradient analysis at the level of individual HMM states, we show that the effect along the principal sensory–transmodal axis (G1) is driven primarily by the states that EBs occupy most frequently, whereas differences along the secondary gradient (G2) are present across all states, with the strongest effect in the state preferentially expressed by controls. These results demonstrate that alterations in functional gradients are themselves state-dependent and tightly coupled to group differences in temporal dynamics. Together, these findings extend previous gradient-based evidence of occipital reorganization in blindness (59, 60) in two important ways: first, they show that the higher segregation from sensorimotor network and integration with transmodal regions is preserved when connectivity is estimated from a generative, model-based framework; and second, they reveal that this repositioning of occipital regions within functional hierarchies is embedded in specific brain states rather than being a static property of the connectome. In the context of crossmodal plasticity, this suggests that the integration of the occipital cortex into transmodal systems is dynamically implemented through the selective recruitment of specific latent states.

Finally, we examined whether early visual deprivation reshapes the low-dimensional geometry of latent brain states. Previous work has shown that large-scale brain dynamics unfold on a low-dimensional manifold aligned with the principal functional gradients in sighted individuals (61, 96, 97). We found that this geometric organization is largely preserved in blindness: in both groups, latent states spanned the principal gradients of cortical organization, suggesting that early visual deprivation leaves the global low-dimensional scaffold of brain dynamics intact while selectively modulating the temporal expression of states within this space. Notably, the state located near the center of the manifold was a low-activation state (state 2), previously described as a transitional hub (Chen et al., 2016; Song et al., 2023), attracting transitions from other brain states. In our study, we confirmed this “transitional” role, as the transition probability analyses revealed that this state attracts transitions from other states regardless of the sensory deprivation. Taken together, these findings reinforce the idea that functional plasticity in blindness operates primarily through changes in temporal dynamics, rather than through a reorganization of the core dynamical architecture.

This study is not exempt from limitations related to the method used. Indeed, the data-driven approach of HMM has an intrinsic limitation represented by the choice of the number of brain states as a priori decision that needs to be made to fit the model. Although we attempted to optimize the process of model order decision and compared results obtained from different model orders, it is important to acknowledge this as a limitation of the study. Furthermore, the concatenation of the timecourses to estimate the parameters implicitly forces the model to estimate the same brain states for all subjects, potentially masking between-group differences in brain states. While the same method could be used to estimate states from the data of each group (blind or sighted), this would not allow for a direct statistical comparison between the metrics of the two groups, as performed in our study. We addressed this limitation by inspecting the Gamma signal at subject level and ensuring that each state is visited by both groups, thus ruling out the possibility that one state clearly describes only one group. Future research could benefit from validating these findings with larger samples or optimized acquisition protocols.

## Conclusions

Our findings highlight the utility of Hidden Markov Models (HMMs) for characterizing the temporal dynamics of functional connectivity and their reorganization following sensory deprivation. We observed that early visual deprivation leads to pronounced alterations in the chronnectome, with brain states dominated by unimodal activity and increased occipito-frontal coupling. Importantly, the hyperconnectivity observed in early blind individuals originates from alterations in the temporal dynamics of brain states, rather than changes in connectivity patterns per se, whereas the patterns of hypoconnectivity arise more broadly across multiple states. Despite these experience-dependent changes, the global low-dimensional organization of brain states and principal functional gradients remains preserved, consistent with a genetically determined cortical scaffold. Together, these results support the idea that robust structural constraints establish the core hierarchy of brain dynamics, while sensory experience drives flexible, state-specific refinements, shaping how the brain dynamically explores its functional repertoire.

## Method

### Participants

In this study we integrated two datasets collected at different times and locations to bolster statistical power. Each dataset comprised EBs and age-and gender-matched sighted controls. The first dataset was obtained in Trento, Italy, in 2023. It consists of forty-eight native Italian speakers, 16 early blind (10 female, mean age=32.8, std=4.6) and 32 sighted (17 female, mean age=30.4, std=7.5). Further details are available in (98). The second dataset, gathered in Montreal, Canada, in 2017, comprised thirty-five subjects, 17 congenitally blind (7 female, mean age=51, std=14.33) and 18 sighted (7 female, mean age = 52.11, std = 12.42). None of the participants had a history of neurological or psychiatric disorders.

### Ethics statement

This study was conducted according to the principles expressed in the Declaration of Helsinki. For the dataset from Trento the ethical committee of the University of Trento approved the experimental protocol in this study (protocol 2014–007). All participants provided written informed consent and were paid for their time. For the dataset from Montreal, all of the procedures were approved by the research ethic and scientific boards of the Centre for Interdisciplinary Research in Rehabilitation of Greater Montreal and the Quebec Bio-Imaging Network. Experiments were undertaken with the understanding and written consent of each subject.

### Data acquisition

The two datasets were acquired using the following protocols:

**Dataset Trento (2023)**. MRI data were collected at the Center for Mind/Brain Sciences, University of Trento (Italy), using a MAGNETOM Prisma 3T MR scanner (Siemens) equipped with a 64-channel head-neck coil. Functional imaging utilized a simultaneous multislice echoplanar imaging sequence, with a scanning plane parallel to the bicommissural plane and a phase encoding direction from anterior to posterior. Parameters included a repetition time (TR) of 1000 ms, echo time (TE) of 28 ms, and a flip angle (FA) of 59°, with a multiband factor of 5, for a total of 480 volumes. For both the early blind and sighted control groups, as well as seven participants in the independent sighted group, images were acquired at a spatial resolution of 3 mm, with a field of view (FOV) of 198 mm × 198 mm, a matrix size of 66 × 66, and 65 axial slices. Slice thickness (ST) was set to 3 mm, with a 0.3 mm gap, resulting in voxel dimensions of 3 × 3 × (3 + 0.3) mm. For the remaining nine participants in the independent sighted group, a higher spatial resolution of 2 mm was employed, with a FOV of 200 mm × 200 mm, a matrix size of 100 × 100, and 65 axial slices. Here, slice thickness was reduced to 2 mm, with a 0.2 mm gap, resulting in voxel dimensions of 2 × 2 × (2 + 0.2) mm. Three-dimensional T1-weighted images were acquired using a magnetization-prepared rapid gradient-echo sequence in the sagittal plane, with parameters including a TR of 2,140 ms, TE of 2.9 ms, inversion time of 950 ms, FA of 12°, FOV of 288 mm × 288 mm, a matrix size of 288 × 288, and 208 continuous sagittal slices. Slice thickness was 1 mm, resulting in voxel dimensions of 1 × 1 × 1 mm.

**Dataset Montreal (2017)**. MRI data were collected in Montreal (Canada), using a Siemens Prisma fit 3T MR scanner equipped with a 32-channel head-neck coil. Functional imaging utilized a simultaneous multislice echoplanar imaging sequence, with a scanning plane parallel to the bicommissural plane and a phase encoding direction from anterior to posterior. Parameters included a repetition time (TR) of 785 ms, echo time (TE) of 3 ms, and a flip angle (FA) of 54°, with a multiband factor of 3, for a total of 472 volumes. For both the early blind and sighted control groups images were acquired at a spatial resolution of 3 mm, with a field of view (FOV) of 192 mm × 192 mm, a matrix size of 64 × 64, and 42 axial slices. Slice thickness (ST) was set to 3 mm, resulting in voxel dimensions of 3 × 3 × 3 mm. Three-dimensional T1-weighted images were acquired using a magnetization-prepared rapid gradient-echo sequence in the sagittal plane, with parameters including a TR of 2,300 ms, TE of 2.26 ms, inversion time of 900 ms, FA of 8°, FOV of 288 mm × 288 mm, a matrix size of 256 × 256, and 176 continuous sagittal slices. Slice thickness was 1 mm, resulting in voxel dimensions of 1 × 1 × 1 mm.

### MRI data preprocessing

The MRI scans were organized into the Brain Imaging Data Structure (BIDS) format using *dcm2niix* (99) and *dcm2bids* (100). Subsequently, the images underwent preprocessing using the fMRIPrep pipeline (101) version 20.2.3. This pipeline included motion correction (*mcflirt*), spatial normalization, co-registration between functional and anatomical scans (*bbregister*), and resampling into a standard space (MNI, 3mm). Following preprocessing, we performed ICA-based denoising (102) at the subject level by manually classifying motion and signal components. Approximately 8% of the components were retained (with 92% classified as noise) in both groups and datasets. The cleaned data were then temporally filtered using a bandpass filter with 0.01 to 0.1 Hz frequency cutoffs to remove noise, enhance the signal-to-noise ratio, and focus on the resting state fluctuations of interest.

To account for potential differences between blind and sighted individuals, we utilized an Atlas parcellation approach instead of conducting a group spatial-ICA. Specifically, we employed the Schaefer Atlas (103), which consists of 100 parcels corresponding to seven resting-state networks (104): visual, somatomotor, dorsal and salience attention, limbic, frontoparietal, and default mode networks. Subsequently, the functional data were coregistered to the Atlas space (MNI, 2mm), and the first eigenvariate was extracted from each parcel using *fslmeants* (*-eig*). This procedure calculates the principal component(s) of the time series within each parcel, providing a single time series that summarizes the dominant pattern of activity while accounting for the highest variance in the parcel’s time series. To equalize the number of volumes across datasets while ensuring signal stabilization and minimizing data loss, we discarded the first 16 volumes (16 s) from the dataset from Trento and the first 8 volumes (6.28 s) from the dataset from Montreal, resulting in 464 volumes per subject.

### Hidden Markov Model

In this study, we adopted a methodology previously outlined in (55), to estimate the temporal dynamic of hidden states by leveraging the HMM-MAR Matlab toolbox (50) publicly available at (https://github.com/OHBA-analysis/HMM-MAR). We utilized the first eigenvariate extracted from each parcel of the Schaefer Atlas (103) for each subject, as input data. Prior to inferring the hidden states, we standardized the subject-specific timecourse of each parcel to have a zero mean and unit standard deviation. Subsequently, we temporally concatenated these standardized values to create a data matrix with dimensions N x V, where N=100 (number of parcels), and V=38512 number of volumes: 83 subjects * 464 volumes). The HMM was then applied to the concatenated data to derive a unified set of brain states shared among all subjects. State inference was performed using a multivariate Gaussian observation model, in which each state is characterized by a state-specific mean activity pattern and functional connectivity (covariance) structure. To ensure robust estimation given the high dimensionality of the data relative to the number of time points, inference was conducted in a reduced low-dimensional space obtained via principal component analysis. Model estimation was repeated multiple times to account for the nonlinearity of the variational Bayes optimization, and the solution with the lowest free energy was selected (see SI for additional details).

The number of hidden states is a critical hyper-parameter that requires a priori specification before initiating the inference process. We explored a range from 2 to 15 states, consistent with prior work (25, 48, 49, 52, 55, 105). Based on model evidence and stability criteria, we selected a solution with 7 states (see SI - *Choice of model size*). Although the inferred states were common across subjects, subject-specific temporal dynamics were obtained using a dual-estimation procedure. This approach yielded individual-level estimates of state time courses, fractional occupancy, transition probability matrices, and state-specific functional connectivity, enabling between-group comparisons of both temporal and spatial features of dynamic functional connectivity.

### Functional connectome over time

The primary aim of this study was to investigate whether early visual deprivation alters the temporal dynamics of functional connectivity. To this end, we compared early blind and sighted individuals across key metrics derived from a Hidden Markov Model (HMM) framework, focusing on subject-level differences in the temporal expression of latent brain states. Group differences were assessed using a GLM-based permutation testing approach while controlling for motion, scanner type, and age, with correction for multiple comparisons. We specifically examined three complementary metrics capturing different aspects of state dynamics: fractional occupancy, reflecting the proportion of time spent in each state; mean lifetime (or dwell time), indexing the duration of individual state visits; and switching rate, measuring the overall stability of state transitions. Together, these measures provide a comprehensive characterization of large-scale brain dynamics in early blind and sighted individuals.

The second aim of the study was to determine whether these alterations in temporal dynamics help explain the differences in stationary functional connectivity reported in blindness. To address this question, we compared large-scale functional connectomes between groups using Network-Based Statistics (NBS), applied to both empirical resting-state FC and model-based FC derived from the HMM. We observed robust group differences in stationary FC, consistent with previous reports of altered occipital connectivity following early visual deprivation. To directly test whether these differences arise from altered temporal dynamics or from intrinsic changes in connectivity structure, we quantified the contribution of HMM-derived brain states to stationary FC. Edge-wise regression analyses revealed that a substantial proportion of variance in stationary FC could be explained by the fractional occupancy–weighted combination of brain states. Crucially, leave-one-state-out reconstructions showed that removing the two states more frequently occupied by early blind participants abolished the characteristic hyperconnectivity pattern, whereas excluding other states had minimal impact. Targeted perturbation analyses further demonstrated that attenuating between-group differences in fractional occupancy but not in state-specific connectivity, was sufficient to eliminate group differences in stationary FC (see SI for additional details).

Together, these analyses indicate that altered temporal expression of a small number of brain states, rather than changes in connectivity structure per se, drives the static FC differences observed in early blindness.

### Functional Connectivity Gradients

Functional gradients provide a low-dimensional representation of the continuous spatial organization of cortical functional connectivity (57). To examine how early visual deprivation impacts this organization, we analyzed functional gradients derived from both model-based stationary connectivity and state-specific connectivity estimated by the HMM. Publicly available reference gradients (NeuroVault collection 1598, https://neurovault.org/collections/1598/) were used to characterize parcel-wise positions in gradient space (see SI for additional details).

We first assessed whether visual deprivation alters the large-scale spatial organization of cortical gradients. Group-level functional connectivity matrices were computed separately for early blind (EB) and sighted control (SC) participants using (i) model-based stationary FC and (ii) state-specific FC patterns. Gradients were derived from these matrices using diffusion map embedding and aligned across groups to the reference gradients using Procrustes rotation. Individual gradient scores were obtained by projecting subject-level connectivity onto the aligned group gradients. We then quantified functional differentiation by measuring gradient range and the separation between the visual network and other resting-state networks along the principal gradients.

Next, we examined how HMM-derived latent brain states are embedded within this low-dimensional gradient space. For each state, mean activation patterns were projected onto the reference gradients, yielding state-specific coordinates for each group. This allowed us to directly compare the spatial positioning of corresponding states between groups and to assess whether latent states occupied distinct and extreme locations along the principal functional axes. To determine whether the observed state organization exceeded that expected by chance, we compared empirical state distributions against surrogate models derived from temporally shifted data.

Together, these analyses allowed us to test whether early visual deprivation alters the spatial embedding of connectivity states within the brain’s principal functional gradients, and whether such changes coexist with a preserved low-dimensional scaffold of large-scale brain organization.

## References

1. T. Elbert, et al., Expansion of the tonotopic area in the auditory cortex of the blind. J. Neurosci. 22, 9941–9944 (2002).

2. I. Hertrich, S. Dietrich, H. Ackermann, Tracking the speech signal--time-locked MEG signals during perception of ultra-fast and moderately fast speech in blind and in sighted listeners. Brain Lang. 124, 9–21 (2013).

3. S. Mattioni, M. Rezk, C. Battal, J. Vadlamudi, O. Collignon, Impact of blindness onset on the representation of sound categories in occipital and temporal cortices. Elife 11 (2022).

4. J. Frasnelli, O. Collignon, P. Voss, F. Lepore, Crossmodal plasticity in sensory loss. Prog. Brain Res. 191, 233–249 (2011).

5. H. Neville, D. Bavelier, Human brain plasticity: evidence from sensory deprivation and altered language experience. Prog. Brain Res. 138, 177–188 (2002).

6. O. Collignon, et al., Functional specialization for auditory–spatial processing in the occipital cortex of congenitally blind humans. Proceedings of the National Academy of Sciences 108, 4435–4440 (2011).

7. G. Dormal, O. Collignon, Functional selectivity in sensory-deprived cortices. J. Neurophysiol. 105, 2627–2630 (2011).

8. E. Ricciardi, G. Handjaras, P. Pietrini, The blind brain: how (lack of) vision shapes the morphological and functional architecture of the human brain. Exp. Biol. Med. (Maywood) 239, 1414–1420 (2014).

9. S. Mattioni, et al., Categorical representation from sound and sight in the ventral occipito-temporal cortex of sighted and blind. Elife 9 (2020).

10. T. R. Makin, J. W. Krakauer, Against cortical reorganisation. Elife 12 (2023).

11. L. Reich, S. Maidenbaum, A. Amedi, The brain as a flexible task machine. Curr. Opin. Neurol. 25, 86–95 (2012).

12. K. J. Friston, L. Harrison, W. Penny, Dynamic causal modelling. Neuroimage 19, 1273–1302 (2003).

13. K. A. Smitha, et al., Resting state fMRI: A review on methods in resting state connectivity analysis and resting state networks. Neuroradiol. J. 30, 305–317 (2017).

14. A. S. Bock, I. Fine, Anatomical and functional plasticity in early blind individuals and the mixture of experts architecture. Front. Hum. Neurosci. 8, 971 (2014).

15. S. Abboud, L. Cohen, Distinctive Interaction Between Cognitive Networks and the Visual Cortex in Early Blind Individuals. Cereb. Cortex 29, 4725–4742 (2019).

16. C. M. Bauer, et al., Multimodal MR-imaging reveals large-scale structural and functional connectivity changes in profound early blindness. PLoS One 12, e0173064 (2017).

17. H. Burton, A. Z. Snyder, M. E. Raichle, Resting state functional connectivity in early blind humans. Front. Syst. Neurosci. 8, 51 (2014).

18. Y. Liu, et al., Whole brain functional connectivity in the early blind. Brain 130, 2085–2096 (2007).

19. E. Striem-Amit, et al., Functional connectivity of visual cortex in the blind follows retinotopic organization principles. Brain 138, 1679–1695 (2015).

20. M. Pelland, et al., State-dependent modulation of functional connectivity in early blind individuals. Neuroimage 147, 532–541 (2017).

21. Z. Wen, et al., Altered functional connectivity of primary visual cortex in late blindness. Neuropsychiatr. Dis. Treat. 14, 3317–3327 (2018).

22. B. Deen, R. Saxe, M. Bedny, Occipital cortex of blind individuals is functionally coupled with executive control areas of frontal cortex. J. Cogn. Neurosci. 27, 1633–1647 (2015).

23. G. Deco, V. K. Jirsa, A. R. McIntosh, Emerging concepts for the dynamical organization of resting-state activity in the brain. Nat. Rev. Neurosci. 12, 43–56 (2011).

24. A. Kucyi, M. J. Hove, M. Esterman, R. M. Hutchison, E. M. Valera, Dynamic brain network correlates of spontaneous fluctuations in attention. Cereb. Cortex 27, 1831–1840 (2017).

25. E. Damaraju, et al., Dynamic functional connectivity analysis reveals transient states of dysconnectivity in schizophrenia. Neuroimage Clin 5, 298–308 (2014).

26. R. M. Hutchison, et al., Dynamic functional connectivity: promise, issues, and interpretations. Neuroimage 80, 360–378 (2013).

27. G. Deco, V. K. Jirsa, Ongoing cortical activity at rest: criticality, multistability, and ghost attractors. J. Neurosci. 32, 3366–3375 (2012).

28. E. C. A. Hansen, D. Battaglia, A. Spiegler, G. Deco, V. K. Jirsa, Functional connectivity dynamics: modeling the switching behavior of the resting state. Neuroimage 105, 525–535 (2015).

29. J. Cabral, M. L. Kringelbach, G. Deco, Exploring the network dynamics underlying brain activity during rest. Prog. Neurobiol. 114, 102–131 (2014).

30. M. Breakspear, Dynamic models of large-scale brain activity. Nat. Neurosci. 20, 340–352 (2017).

31. T. T. Nakagawa, V. K. Jirsa, A. Spiegler, A. R. McIntosh, G. Deco, Bottom up modeling of the connectome: linking structure and function in the resting brain and their changes in aging. Neuroimage 80, 318–329 (2013).

32. M. A. Gainey, D. E. Feldman, Multiple shared mechanisms for homeostatic plasticity in rodent somatosensory and visual cortex. Philos. Trans. R. Soc. Lond. B Biol. Sci. 372 (2017).

33. D. Muret, T. R. Makin, The homeostatic homunculus: rethinking deprivation-triggered reorganisation. Curr. Opin. Neurobiol. 67, 115–122 (2021).

34. A. Kral, P. A. Yusuf, R. Land, Higher-order auditory areas in congenital deafness: Top-down interactions and corticocortical decoupling. Hear. Res. 343, 50–63 (2017).

35. C. Lubinus, et al., Data-Driven Classification of Spectral Profiles Reveals Brain Region-Specific Plasticity in Blindness. Cereb. Cortex 31, 2505–2522 (2021).

36. R. Pant, et al., Stimulus-evoked and resting-state alpha oscillations show a linked dependence on patterned visual experience for development. Neuroimage Clin 38, 103375 (2023).

37. A. Kriegseis, E. Hennighausen, F. Rösler, B. Röder, Reduced EEG alpha activity over parieto-occipital brain areas in congenitally blind adults. Clin. Neurophysiol. 117, 1560–1573 (2006).

38. I. M. Schepers, J. F. Hipp, T. R. Schneider, B. Röder, A. K. Engel, Functionally specific oscillatory activity correlates between visual and auditory cortex in the blind. Brain 135, 922–934 (2012).

39. D. J. Hawellek, et al., Altered intrinsic neuronal interactions in the visual cortex of the blind. J. Neurosci. 33, 17072–17080 (2013).

40. K. Rączy, et al., Typical resting-state activity of the brain requires visual input during an early sensitive period. Brain Commun 4, fcac146 (2022).

41. S. S. Menon, K. Krishnamurthy, A Comparison of Static and Dynamic Functional Connectivities for Identifying Subjects and Biological Sex Using Intrinsic Individual Brain Connectivity. Sci. Rep. 9, 5729 (2019).

42. A. D. Savva, G. D. Mitsis, G. K. Matsopoulos, Assessment of dynamic functional connectivity in resting-state fMRI using the sliding window technique. Brain Behav. 9, e01255 (2019).

43. E. W. S. Carbo, et al., Dynamic hub load predicts cognitive decline after resective neurosurgery. Sci. Rep. 7, 42117 (2017).

44. J. Kim, et al., Abnormal intrinsic brain functional network dynamics in Parkinson’s disease. Brain 140, 2955–2967 (2017).

45. T. Watanabe, G. Rees, Brain network dynamics in high-functioning individuals with autism. Nat. Commun. 8, 16048 (2017).

46. A. J. Quinn, et al., Task-Evoked Dynamic Network Analysis Through Hidden Markov Modeling. Front. Neurosci. 12, 603 (2018).

47. J. Taghia, et al., Uncovering hidden brain state dynamics that regulate performance and decision-making during cognition. Nat. Commun. 9, 2505 (2018).

48. M. Moretto, E. Silvestri, A. Zangrossi, M. Corbetta, A. Bertoldo, Unveiling whole-brain dynamics in normal aging through Hidden Markov Models. Hum. Brain Mapp. 43, 1129–1144 (2022).

49. M. Moretto, et al., The dynamic functional connectivity fingerprint of high-grade gliomas. Sci. Rep. 13, 10389 (2023).

50. D. Vidaurre, et al., Spectrally resolved fast transient brain states in electrophysiological data. Neuroimage 126, 81–95 (2016).

51. B. Cao, et al., Abnormal dynamic properties of functional connectivity in disorders of consciousness. Neuroimage Clin 24, 102071 (2019).

52. A. Kottaram, et al., Brain network dynamics in schizophrenia: Reduced dynamism of the default mode network. Hum. Brain Mapp. 40, 2212–2228 (2019).

53. J. Van Schependom, et al., Altered transient brain dynamics in multiple sclerosis: Treatment or pathology? Hum. Brain Mapp. 40, 4789–4800 (2019).

54. T. A. W. Bolton, E. Morgenroth, M. G. Preti, D. Van De Ville, Tapping into multi-faceted human behavior and psychopathology using fMRI brain dynamics. Trends Neurosci. 43, 667–680 (2020).

55. D. Vidaurre, S. M. Smith, M. W. Woolrich, Brain network dynamics are hierarchically organized in time. Proc. Natl. Acad. Sci. U. S. A. 114, 12827–12832 (2017).

56. A. I. Luppi, et al., Unravelling consciousness and brain function through the lens of time, space, and information. Trends Neurosci. 47, 551–568 (2024).

57. D. S. Margulies, et al., Situating the default-mode network along a principal gradient of macroscale cortical organization. Proc. Natl. Acad. Sci. U. S. A. 113, 12574–12579 (2016).

58. J. M. Huntenburg, P.-L. Bazin, D. S. Margulies, Large-scale gradients in human cortical organization. Trends Cogn. Sci. 22, 21–31 (2018).

59. M. Tamietto, et al., Innate cortical gradients constrain cross-modal plasticity. Research Square (2025).

60. C. Koba, et al., Visual experience shapes functional connectome gradients. bioRxiv (2025).

61. H. Song, W. M. Shim, M. D. Rosenberg, Large-scale neural dynamics in a shared low-dimensional state space reflect cognitive and attentional dynamics. Elife 12 (2023).

62. S. Chen, J. Langley, X. Chen, X. Hu, Spatiotemporal modeling of brain dynamics using resting-state functional magnetic resonance imaging with Gaussian hidden Markov model. Brain Connect. 6, 326–334 (2016).

63. M. J. S. Guerreiro, M. Linke, S. Lingareddy, R. Kekunnaya, B. Röder, The effect of congenital blindness on resting-state functional connectivity revisited. Sci. Rep. 11, 1–14 (2021).

64. C. Koba, A. Crimi, O. Collignon, E. Ricciardi, U. Hasson, Neural networks associated with eye movements in congenital blindness. Eur. J. Neurosci. (2024). 10.1111/ejn.16459.

65. M. Tian, X. Xiao, H. Hu, R. Cusack, M. Bedny, Visual experience shapes functional connectivity between occipital and non-visual networks. (2024).

66. B. Vázquez-Rodríguez, et al., Gradients of structure-function tethering across neocortex. Proc. Natl. Acad. Sci. U. S. A. 116, 21219–21227 (2019).

67. M. G. Preti, D. Van De Ville, Decoupling of brain function from structure reveals regional behavioral specialization in humans. Nat. Commun. 10, 4747 (2019).

68. C. Paquola, et al., Shifts in myeloarchitecture characterise adolescent development of cortical gradients. Elife 8 (2019).

69. J. B. Burt, et al., Hierarchy of transcriptomic specialization across human cortex captured by structural neuroimaging topography. Nat. Neurosci. 21, 1251–1259 (2018).

70. L. Ortiz-Terán, et al., Brain Plasticity in Blind Subjects Centralizes Beyond the Modal Cortices. Front. Syst. Neurosci. 10, 61 (2016).

71. A. Amedi, N. Raz, P. Pianka, R. Malach, E. Zohary, Early “visual” cortex activation correlates with superior verbal memory performance in the blind. Nat. Neurosci. 6, 758–766 (2003).

72. A. Amedi, A. Floel, S. Knecht, E. Zohary, L. G. Cohen, Transcranial magnetic stimulation of the occipital pole interferes with verbal processing in blind subjects. Nat. Neurosci. 7, 1266–1270 (2004).

73. M. Bedny, A. Pascual-Leone, D. Dodell-Feder, E. Fedorenko, R. Saxe, Language processing in the occipital cortex of congenitally blind adults. Proc. Natl. Acad. Sci. U. S. A. 108, 4429–4434 (2011).

74. O. Collignon, et al., Impact of blindness onset on the functional organization and the connectivity of the occipital cortex. Brain 136, 2769–2783 (2013).

75. L. Gagnon, R. Kupers, M. Ptito, Neural correlates of taste perception in congenital blindness. Neuropsychologia 70, 227–234 (2015).

76. F. Gougoux, et al., Voice perception in blind persons: a functional magnetic resonance imaging study. Neuropsychologia 47, 2967–2974 (2009).

77. N. Raz, A. Amedi, E. Zohary, V1 activation in congenitally blind humans is associated with episodic retrieval. Cereb. Cortex 15, 1459–1468 (2005).

78. U. Noppeney, The effects of visual deprivation on functional and structural organization of the human brain. Neurosci. Biobehav. Rev. 31, 1169–1180 (2007).

79. D. Radziun, M. Korczyk, L. Crucianelli, M. Szwed, H. H. Ehrsson, Heartbeat counting accuracy is enhanced in blind individuals. J. Exp. Psychol. Gen. 152, 2026–2039 (2023).

80. D. Radziun, L. Crucianelli, M. Korczyk, M. Szwed, H. H. Ehrsson, The perception of affective and discriminative touch in blind individuals. Behav. Brain Res. 444, 114361 (2023).

81. D. Goldreich, I. M. Kanics, Performance of blind and sighted humans on a tactile grating detection task. Percept. Psychophys. 68, 1363–1371 (2006).

82. C. Y. Wan, A. G. Wood, D. C. Reutens, S. J. Wilson, Congenital blindness leads to enhanced vibrotactile perception. Neuropsychologia 48, 631–635 (2010).

83. C. Battal, V. Occelli, G. Bertonati, F. Falagiarda, O. Collignon, General enhancement of spatial hearing in congenitally blind people. Psychol. Sci. 31, 1129–1139 (2020).

84. M. Bedny, Evidence from Blindness for a Cognitively Pluripotent Cortex. Trends Cogn. Sci. 21, 637–648 (2017).

85. J. Duncan, The multiple-demand (MD) system of the primate brain: mental programs for intelligent behaviour. Trends Cogn. Sci. 14, 172–179 (2010).

86. M. D. Fox, M. E. Raichle, Spontaneous fluctuations in brain activity observed with functional magnetic resonance imaging. Nat. Rev. Neurosci. 8, 700–711 (2007).

87. Z. Zhang, et al., Intrinsic neural linkage between primary visual area and default mode network in human brain: Evidence from visual mental imagery. Neuroscience 379, 13–21 (2018).

88. J. Bergmann, E. Genç, A. Kohler, W. Singer, J. Pearson, Smaller primary visual cortex is associated with stronger, but less precise mental imagery. Cereb. Cortex 26, 3838–3850 (2016).

89. S. M. Daselaar, Y. Porat, W. Huijbers, C. M. A. Pennartz, Modality-specific and modality-independent components of the human imagery system. Neuroimage 52, 677–685 (2010).

90. A. Amedi, R. Malach, A. Pascual-Leone, Negative BOLD differentiates visual imagery and perception. Neuron 48, 859–872 (2005).

91. G. Iurilli, et al., Sound-driven synaptic inhibition in primary visual cortex. Neuron 73, 814–828 (2012).

92. P. J. Laurienti, et al., Deactivation of sensory-specific cortex by cross-modal stimuli. J. Cogn. Neurosci. 14, 420–429 (2002).

93. M. T. deBettencourt, J. D. Cohen, R. F. Lee, K. A. Norman, N. B. Turk-Browne, Closed-loop training of attention with real-time brain imaging. Nat. Neurosci. 18, 470–475 (2015).

94. F. Setti, et al., A modality-independent proto-organization of human multisensory areas. Nat Hum Behav 7, 397–410 (2023).

95. E. Ricciardi, D. Bottari, M. Ptito, B. Röder, P. Pietrini, The sensory-deprived brain as a unique tool to understand brain development and function. Neurosci. Biobehav. Rev. 108, 78–82 (2020).

96. A. Turnbull, et al., Reductions in task positive neural systems occur with the passage of time and are associated with changes in ongoing thought. Sci. Rep. 10, 9912 (2020).

97. T. Karapanagiotidis, et al., The psychological correlates of distinct neural states occurring during wakeful rest. Sci. Rep. 10, 21121 (2020).

98. Y. Xu, et al., Similar object shape representation encoded in the inferolateral occipitotemporal cortex of sighted and early blind people. PLoS Biol. 21, e3001930 (2023).

99. X. Li, P. S. Morgan, J. Ashburner, J. Smith, C. Rorden, The first step for neuroimaging data analysis: DICOM to NIfTI conversion. J. Neurosci. Methods 264, 47–56 (2016).

100. K. J. Gorgolewski, et al., BIDS apps: Improving ease of use, accessibility, and reproducibility of neuroimaging data analysis methods. PLoS Comput. Biol. 13, e1005209 (2017).

101. O. Esteban, et al., fMRIPrep: a robust preprocessing pipeline for functional MRI. Nat. Methods 16, 111–116 (2019).

102. L. Griffanti, et al., ICA-based artefact removal and accelerated fMRI acquisition for improved resting state network imaging. Neuroimage 95, 232–247 (2014).

103. A. Schaefer, et al., Local-Global Parcellation of the Human Cerebral Cortex from Intrinsic Functional Connectivity MRI. Cereb. Cortex 28, 3095–3114 (2018).

104. B. T. T. Yeo, et al., The organization of the human cerebral cortex estimated by intrinsic functional connectivity. J. Neurophysiol. 106, 1125–1165 (2011).

105. A. P. Baker, et al., Fast transient networks in spontaneous human brain activity. Elife 3, e01867 (2014).

