## Supplementary Material and Figures for "Reorganization of large-scale network dynamics after early visual deprivation"

#### HMM - Choice of the model size

From each model, we extracted metrics such as Free Energy and Average Log-Likelihood (avLL). However, the Free Energy displayed a tendency to decrease with the increase in the number of states without exhibiting any local minima, rendering it unreliable as a criterion for selecting the model's number of states (see Fig. S1). Similarly, the avLL consistently assumed negative values with a monotonic trend, making it challenging to discern a maximum peak. Although we computed the Akaike Information Criterion (AIC) and Bayesian Information Criterion (BIC), they invariably favored the lowest order ($K$=2) as the optimal choice, potentially influenced by the negative avLL values. Given these limitations, we sought alternative measures to inform our decision-making process. First of all, we visually inspected the within-subject dynamics of hidden states to assess whether, in the specific model order, we could extract the single-subject temporal dynamic of hidden states. To do that, for each subject we plotted multiple measures estimated by the HMM: 1). Gamma, that is the time series of probability of state activation as a function of time; 2). the Viterbi path, that is the vector showing the most probable a posteriori state and 3). the Transition Probability Matrix, that is the matrix with the probability of transitioning from one specific state to another (see Fig. S2A for data of a representative subject). We observed that, within the 5-9 state range, all subjects exhibited temporal dynamics of functional connectivity. However, beyond this point, we noticed reduced temporal dynamics mainly in two sighted individuals, belonging to the dataset from Montreal, forcing them to be excluded from the analysis (see Fig. S3). Thus, we chose the range 5-9 as the suitable range for model order. Then, we extracted the ratio of coefficient of variation (CV) of the estimates for each model (1, 2), to quantify the reliability of the mean estimates as the model order increased. Finally, we explored the distribution of mean activations and distances between pairs of functional connectivity matrices, by means of geodesic distance and Pearson dissimilarity (3), hypothesizing that lower correlations and greater distances would indicate more independent states (see Fig. S2B). Our comprehensive analysis pointed to 7 as a strong candidate for the chosen model order. Indeed, within the chosen range, Model 7 displayed the lowest CV ratios (with a ratio of CV=0.12), as well as the most favorable combination of low correlation between mean activations and high distance between FC matrices. Thus, we decided to proceed with 7 as the chosen model order. Nonetheless, we conducted analyses for all orders within the 5-9 range and compared mean activation maps to ensure consistency across model orders.

#### HMM - Inference

The HMM serves as a probabilistic framework that applies variational Bayes inference to extract an approximation of the model posterior distribution of hidden states ($K$) inputting the observable data, i.e. each parcel's timecourse. Each of these hidden states has a certain probability distribution over time and represents a distinct spatiotemporal pattern of activation.
In this study we used a multivariate Gaussian observation model, that specifies a Gaussian distribution of the fMRI data, wherein each state is represented by a specific mean activity and covariance matrix. Given the high number of parcels included (100, atlas based) and the reduced number of timepoints (464 volumes), we opted to estimate the hidden states using a Gaussian observational model within a low-dimensional space (PCA space), retaining 10 components (4). When we attempted to infer the hidden states without any dimensionality reduction, we encountered a lack of discernible single-subject temporal dynamics. This implies that each subject was merely described by a single state persisting over time. The parameters fitted to the HMM on reduced data (PCA) encompass: 1) a vector of the state-specific mean activity for each parcel, when the state $K$ is active (options.zeromean = 0) ; 2) the variance-covariance matrices of N x N, where N represents the number of parcels (100), when each state is active (options.covtype = 'full'); 3) the probabilities of transitioning from one state to another, at any time point.  For initialization, we employed a mixture of Gaussian models with 10 optimization cycles and 3 repetitions. From these repetitions, the one exhibiting the most favorable Free Energy (FE) was selected as the starting point for the inference run. Recognizing the nonlinearity inherent in the Variational Bayes algorithm, we repeated the estimation process 20 times. Subsequently, we ranked these repetitions based on FE, ultimately selecting the one with the lowest value.

#### Functional connectome over time

The primary aim of this study is to investigate disparities in the temporal dynamics of functional connectivity between early blind and sighted individuals. To achieve this, we conducted analyses on three key metrics defining the "Chronnectome", employing a GLM-based permutation testing approach (*hmmtest* function within HMM-MAR toolbox) adjusted to include the mean lifetime, without including the transition probability matrices in the analysis. Our design matrix incorporated the fixed-effect of interest, namely, the early blind vs sighted grouping variable, and three covariates of no interest: mean framewise displacement to account for motion artifacts (a single value per subject), scanner type and age. To address multiple comparisons, we applied the Benjamini–Hochberg false discovery rate procedure, adopting a conservative threshold of $q < 0.025$. Between-group differences were evaluated based on *Gamma,* which estimates the probability of each state to be active over time, at the subject level. This probability is used to extract the three metrics under comparison: (1) fractional occupancy, representing the average state probability across time for each subject and yielding an N x K matrix, where N is the number of subjects (83) and K is the number of states (7); (2) mean lifetime or dwell time, indicating the number of time points spent per visit to a given state; and (3) switching time, which represents the mean frequency of switching between one state and another and serves as a measure of stability (one value per subject). These analyses collectively offer insights into the dynamic temporal patterns characterizing the neural dynamics of early blind and sighted individuals.

The second aim of this study is to determine whether the alterations in temporal dynamics offer insights into differences observed in stationary functional connectomes. For this reason, we investigated large-scale topographical disparities in the functional connectome derived from blind and sighted individuals, by using the Network-Based Statistic Toolbox (NBS) in MATLAB, as outlined by (5). NBS represents a nonparametric statistical approach that extends the cluster-based method used in mass univariate hypothesis testing from physical to topological space, where a cluster's equivalent is a graph component. This allows for the identification of interconnected regions exhibiting significant between-group differences in functional connectivity, providing insights into the network-level between-group differences.

First of all, we quantified stationary functional connectome (FC) using two approaches: (1) the empirical stationary FC, and (2) the time-averaged, model-based FC estimated from the HMM. In the former, we computed the stationary functional connectome by calculating the Pearson correlation between the first eigenvariate of each pair of ROIs. In the latter, we computed a weighted average of the normalized correlation matrices for each state ($100 \times 100$ per subject), with weights given by the fractional occupancies. The fractional occupancy reflects the proportion of time each subject spends in a given state (6). This analysis leads to time-averaged, model-based functional connectome for each subject, that is the functional connectivity estimated by the model, that remains stable over time. In addition, we used the subject-specific HMM, to derive functional connectivity matrices at the individual level, resulting in N x N matrices where N represents the number of parcels (100) when each state K is active. To facilitate group-level comparisons, all these single-subject matrices underwent Fisher's r-to-z transformation. The design matrix used in the NBS analysis integrates the fixed effect of the group alongside mean framewise displacement, scanner type and age as covariates of no interest, mirroring our previous design matrix. We set the primary statistical threshold to 4 to identify supra-threshold connections, and we executed 10,000 permutations to ensure robust statistical inference. Controlling for family-wise error rate at 5%, we scrutinized the resultant network to elucidate significant between-group differences in the total number of connections (*extent*) across the brain's parcellated regions.

To determine whether group differences in stationary functional connectivity (FC) arise from altered temporal dynamics or from intrinsic changes in connectivity patterns, we quantified the contribution of HMM-derived brain states to stationary FC. We performed an edge-wise regression analysis to quantify the contribution of the FC associated with each HMM state to stationary functional connectivity. For each subject, we extracted either the model-based time-averaged FC or the empirical stationary FC. The upper-triangular elements of each FC matrix were vectorized to obtain edge-wise values. For each edge, we fitted a least-squares regression model that included the FO-weighted FC of all HMM states as predictors, along with age, scanner type, and motion as covariates of no interest. To summarize the overall contribution of HMM states to stationary FC, we extracted the adjusted $R^2$ and compared predicted versus observed FC. To assess statistical significance, we performed a permutation test in which the rows of the dynamic predictors were randomly shuffled across subjects. False discovery rate correction was applied to identify significant edges. This analysis allowed us to estimate how much of the structure in both empirical and model-based stationary FC can be explained by the combined influence of all HMM states, while accounting for relevant covariates.

**State-wise contribution to stationary functional connectivity**

To assess whether individual HMM states were necessary for the observed group differences in stationary functional connectivity (FC), we performed a leave-one-state-out reconstruction of subject-level FC matrices. For each subject, alternative FC matrices were reconstructed by excluding the contribution of one state at a time, while preserving the original fractional occupancies of all remaining states. This procedure removes the contribution of the excluded state without redistributing its temporal weight to other states, thereby allowing us to test whether the presence of that state is required to reproduce the group-level differences. Because fractional occupancies were not renormalized after state exclusion, this analysis specifically tests the necessity of each state’s absolute contribution, rather than potential compensatory effects of the remaining states. Network-based statistics (NBS) were then applied to each leave-one-state-out FC matrix using the same design matrix, contrasts, and statistical thresholds as in the full model analysis.

**Disentangling temporal and spatial contributions of HMM states**

Each brain state is characterized by two key components that may differently determine its contribution to the stationary FC profile: **(1)** the state-specific functional connectome and **(2)** chronnectomic metrics that describe the temporal dynamics of the state. Among the metrics extracted, we focused on fractional occupancy, as this metric exhibited significant between-group differences. To assess which features of the HMM states contribute most to the observed between-group differences in stationary FC (empirical or model-based), we independently quantified the contributions of fractional occupancy (FO) and state-specific functional connectivity (FC). To selectively attenuate between-group differences in fractional occupancy (FO) or state-specific functional connectivity (FC) for a given HMM state, we applied a directional reassignment procedure. For each permutation (500 iterations), FO or FC values from one group were randomly reassigned to the other group using sampling with replacement, while the remaining group remained unchanged. Depending on the state of interest, either (i) FO or FC values from control participants were sampled and assigned to early blind (EB) participants (states 1 and 6), or (ii) FO or FC values from EB participants were sampled and assigned to controls (state 7). This preserves the original sample size and marginal distribution of FO or FC values, while selectively disrupting their association with group membership for the targeted state. For each permutation, participant-specific stationary FC matrices were reconstructed as FO-weighted sums of state-specific FC matrices. In the FO reassignment analysis, FO values for the shuffled state were replaced according to the permutation while FO values for all other states remained unchanged. In the FC reassignment analysis, state-specific FC matrices were reassigned according to the permutation while individual FO profiles were kept fixed. Reconstructed FC matrices were Fisher z-transformed to normalize correlation values and served as input to the Network-Based Statistic (NBS) toolbox to test for between-group differences in stationary FC under attenuated FO or FC conditions. As an additional control, we also replaced FO or FC values of the target state with the group mean instead of random reassignment, allowing us to test whether the group-level average alone was sufficient to reproduce between-group differences while removing individual variability. This complementary perturbation framework enables assessment of whether between-group differences in stationary functional connectivity arise primarily from differences in the temporal expression of specific HMM states (FO), from differences in their intrinsic connectivity patterns (FC), or from a combination of both.

#### Functional connectivity gradients

Functional gradients (7) describe continuous axes of functional organization derived from whole-brain connectivity patterns. We obtained the publicly available voxelwise gradient maps (NeuroVault collection 1598, https://neurovault.org/collections/1598/) and averaged gradient values within each of 100 cortical parcels to characterize parcel-wise positions in gradient space.

The first goal was to assess how visual deprivation alters the spatial organization of cortical gradients. To this end, we computed group-level FC matrices separately for EBs and SCs, for (i) the model-based stationary FC and (ii) the FC patterns associated with each hidden Markov model (HMM) state. Each group-level FC matrix was thresholded row-wise to retain the top 10% of strongest connections, ensuring equal sparsity across regions, and used to derive a normalized cosine similarity affinity matrix quantifying the similarity of connectivity profiles between regions. This affinity matrix was embedded into a low-dimensional space using diffusion map embedding, as implemented in the BrainSpace toolbox. To enable comparisons across groups, the resulting gradients were aligned to the reference gradients of Margulies et al. (2016) via Procrustes rotation. Subject-specific gradient scores for each cortical region were then obtained by projecting each individual's unthresholded FC matrix - whether from the stationary FC or from the FC of each HMM state - onto the corresponding aligned group-level gradients. To assess whether visual deprivation alters the degree of segregation between connectivity profiles, we quantified gradient dispersion and inter-network separation. First, for each subject, we computed the range of each gradient as the difference between the maximum and minimum gradient scores across cortical regions (8). Second, we measured the separation between the visual resting-state network (RSN) and all other RSNs along the principal gradients. Specifically, each network was represented by the mean gradient value across its vertices, and the distance between the visual network and each other RSN was defined as the absolute difference between these mean values. Group differences in both measures were assessed using a Wilcoxon rank-sum test. Effect size was quantified using the rank-biserial correlation ($r$), computed from the $z$ statistic of the Wilcoxon rank-sum test ($r = z / \sqrt{N}$), which measures the strength of separation between the two groups' rank distributions. All statistical comparisons were corrected for multiple comparisons using FDR ($q < 0.025$).

The second goal was to map our latent HMM-derived states onto established gradient axes for each group in order to examine how early visual deprivation affects the low-dimensional geometry of latent neural states. To do that, we projected the vector of mean activation for each subject, ROI and state, (*getMean.m)* into the reference gradients. Group-level state coordinates were computed as the mean across subjects within each group. To quantify group differences at the state level, we calculated the Euclidean distance between corresponding states in EB and SC participants. This provided a state-wise measure of divergence in gradient space. To evaluate whether our HMM-derived latent states occupied distinct and extreme positions in the gradient space, we followed the procedure described in (9). As in the original paper, a null distribution was generated using 1,000 surrogate HMMs fitted to parcel-wise circularly time-shifted data, preserving temporal structure while disrupting inter-parcel covariance. Latent state positions were projected onto the same gradient axes, and pairwise Euclidean distances between states were computed. The mean distance between empirical states was compared against the null distribution using a two-tailed permutation test. To assess extremity, empirical state positions along each gradient were compared to surrogate distributions. Statistical significance was corrected for multiple comparisons using false discovery rate (FDR).

### Figures

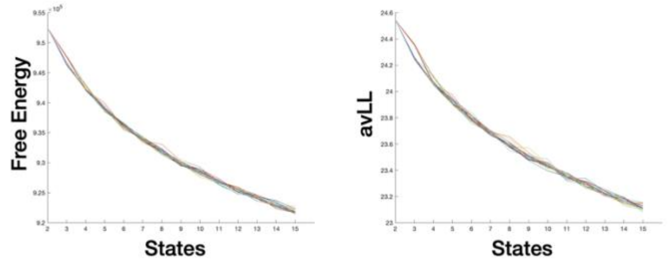


**Figure S1. Choice of the model size.** Free energy and absolute average log-likelihood as a function of the repetition (multiple colors) and the model order, from 2 to 15.

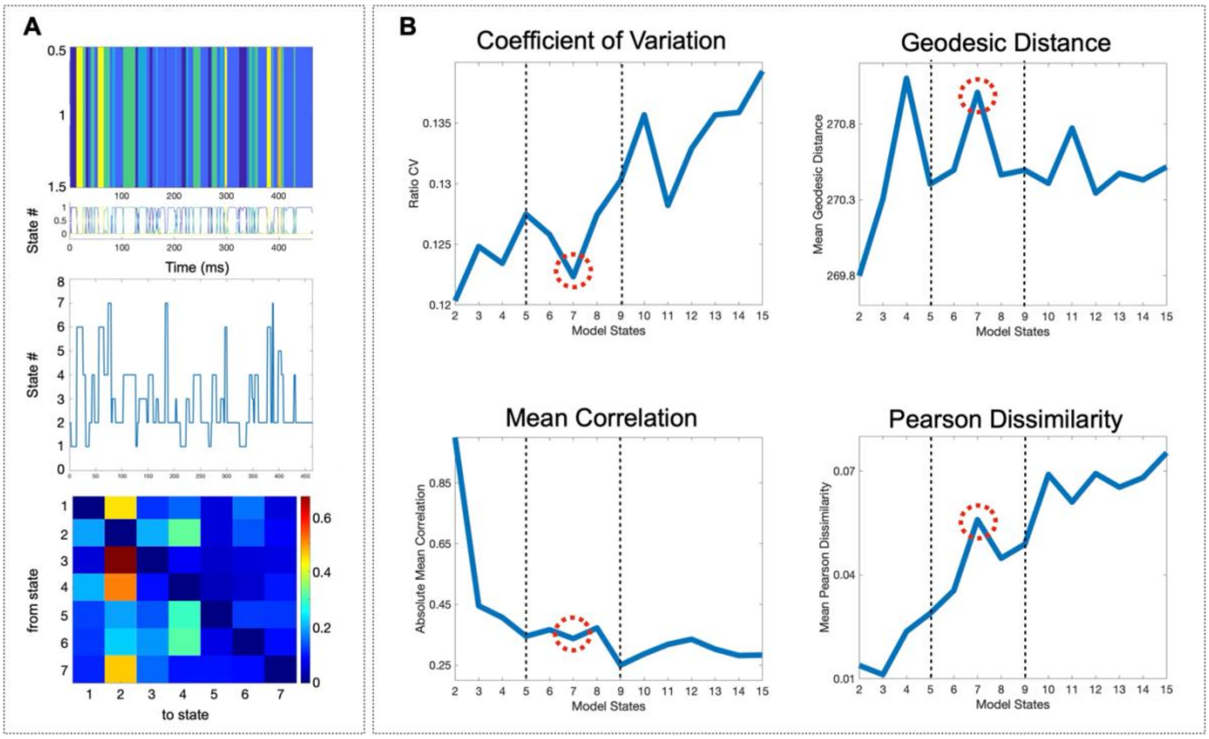


**Figure S2. Choice of the model size.** Visualization of the criteria used to select the order of the HMM. **A.** Visualization of Gamma (upper), Viterbi Path (middle) and transition probability (bottom) for one subject from Dataset 2. **B.** Representation of multiple measures extracted from each model order: Coefficient of Variation, as a measure of reliability of mean estimates as the model order increases; Mean Correlation, Geodesic Distance and Pearson Dissimilarity as measures of independence of hidden states. Lower correlation and higher distance between functional connectivity matrices highlight the presence of more independent states. The dashed red circle indicates the chosen model order.

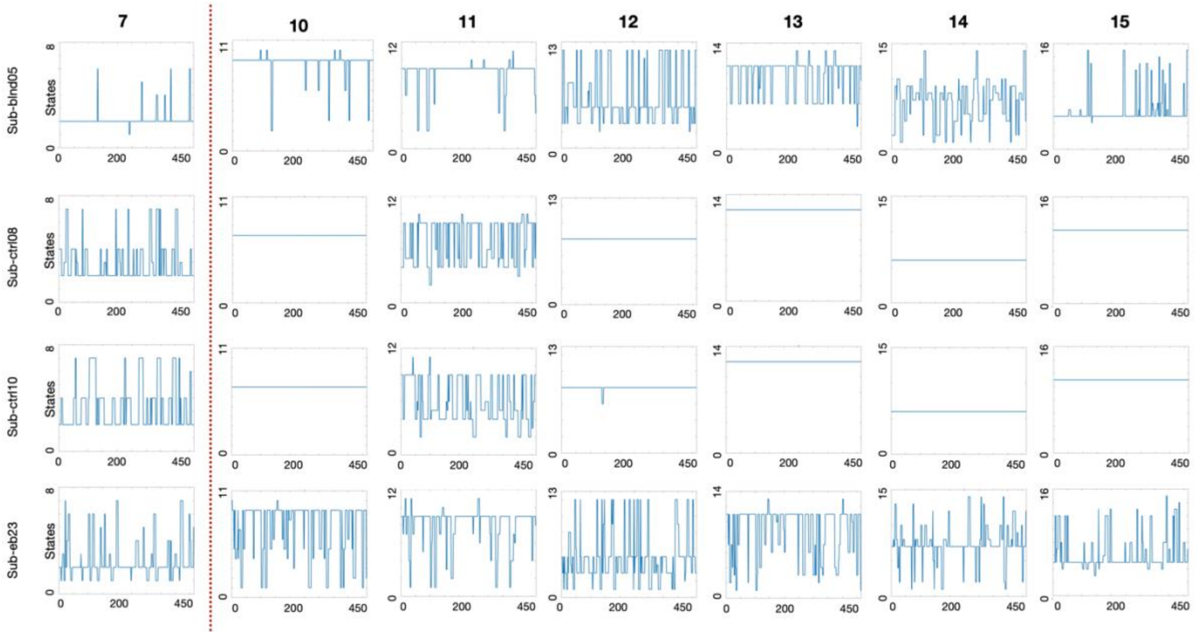


**Figure S3. Choice of the model size.** Visualization of the Viterbi path as a function of time for those subjects who show a reduced temporal dynamic. The increase in the order of the model determines a reduction in the temporal dynamics of a few subjects, mainly belonging to the dataset from Montreal (2017).

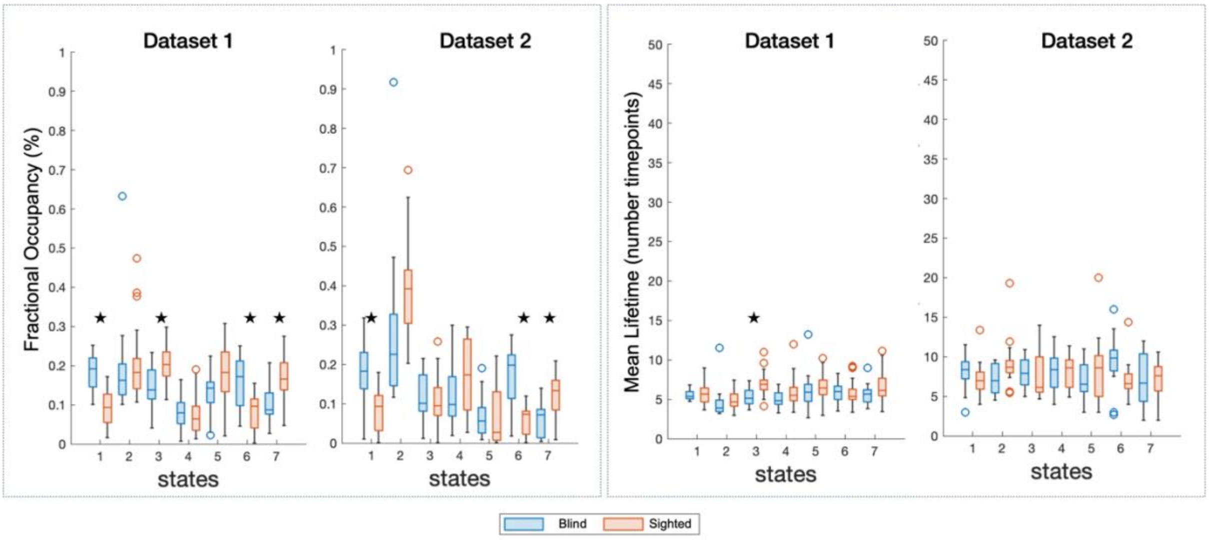


**Figure S4**. Differences in the "chronnectome" between blind and sighted individuals including one dataset. Left: Fractional Occupancy for each group, as a function of state, for the dataset from Trento (left) and the dataset from Montreal (right). Right: Mean Lifetime for each group, as a function of state, for the dataset from Trento (left) and the dataset from Montreal (right).

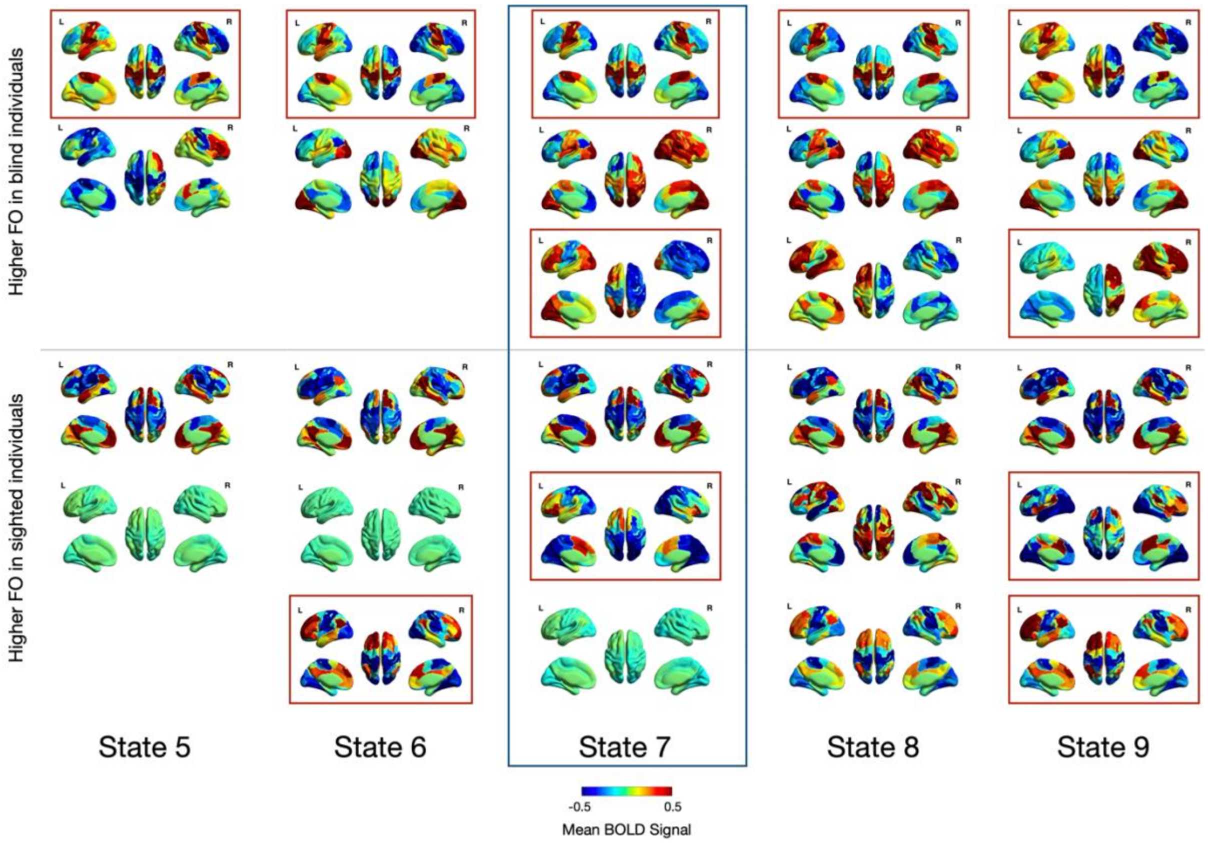


**Figure S5. Maps of mean activation estimated from HMM using different numbers of states (from 5 to 9).** Red squares indicate significant results. The blue square indicates the chosen model order.

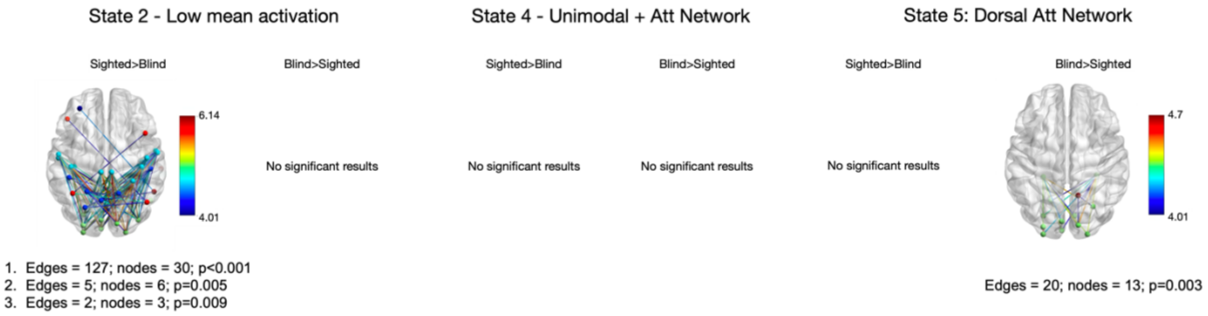


**Figure S6.** Network-level between-group differences in the functional connectome for the time-varying FC of states 2, 4 and 5.

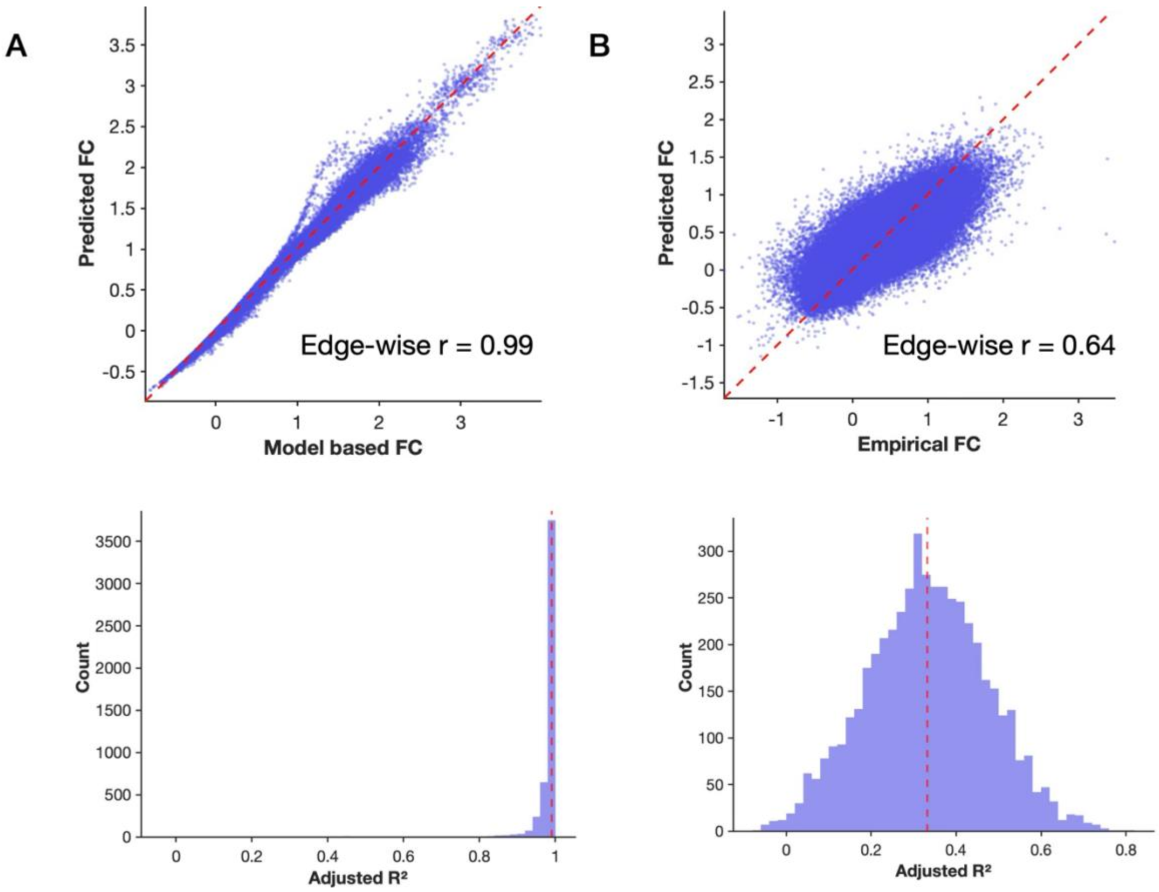


**Fig. S7. Contribution of each HMM-derived state to stationary functional connectivity (FC). A.** Model-based time-averaged stationary FC. **B.** Empirical stationary FC. Upper row: Relationship between model-based time-averaged FC (A) or empirical FC (B) and the HMM-predicted FC. Each point represents one edge. The dashed line represents the identity line. The close clustering around the identity line indicates that dynamic HMM states provide a meaningful approximation of stationary FC. Bottom line: Distribution of the adjusted R2 of the model. The dashed line represents the median adjusted R2.

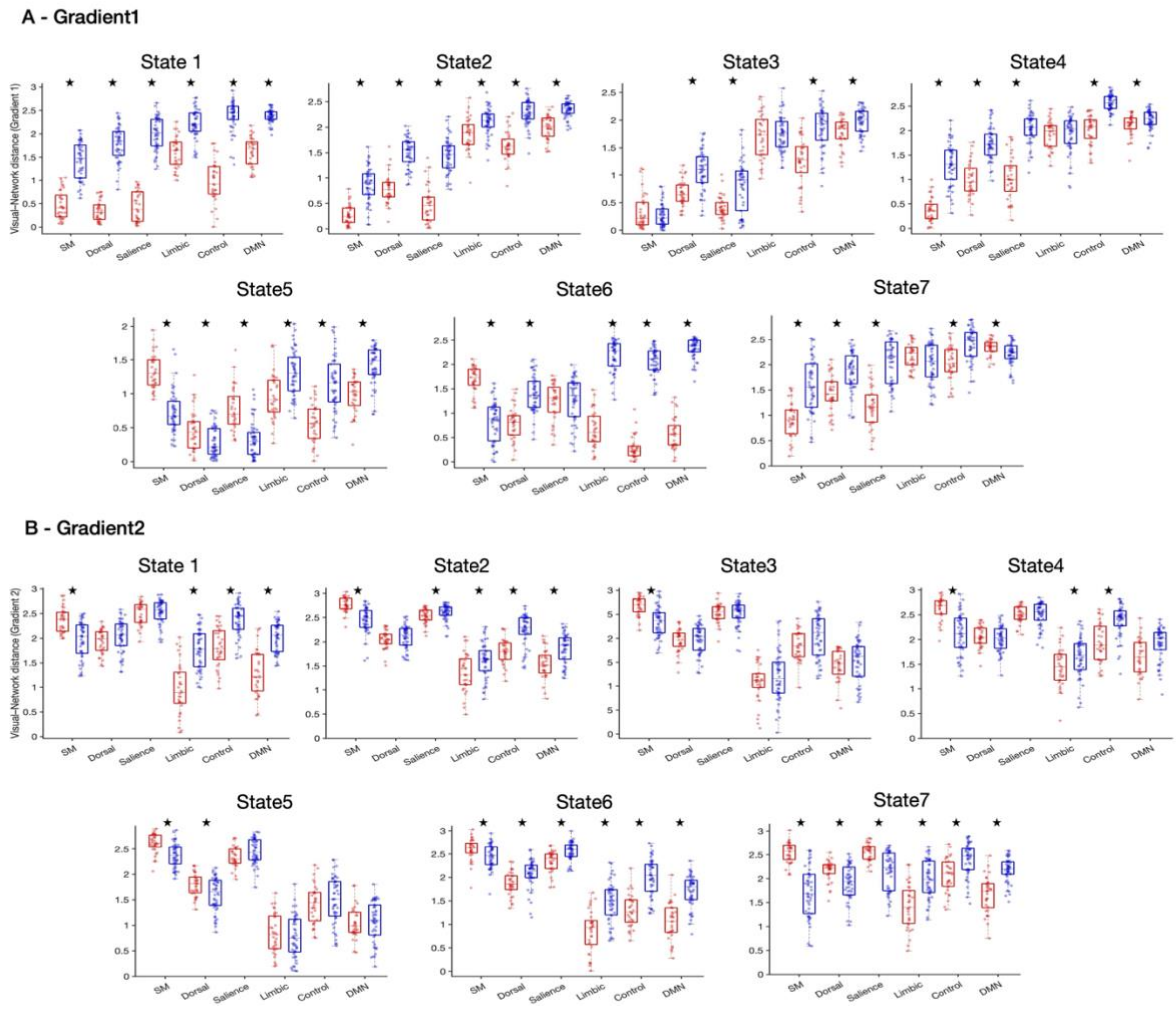


**Fig. S8**. **Absolute Distance between the visual network and all other resting-state networks (RSNs)** for EBs (red) and SCs (blue), shown separately for Gradient 1 (**A**) and Gradient 2 (**B**), for each HMM-state. Asterisks indicate the significant results after correcting for multiple comparisons (FDR).

### References


1. M. Moretto, E. Silvestri, A. Zangrossi, M. Corbetta, A. Bertoldo, Unveiling whole-brain dynamics in normal aging through Hidden Markov Models. *Hum. Brain Mapp.* **43**, 1129–1144 (2022).

2. M. Moretto, *et al.*, The dynamic functional connectivity fingerprint of high-grade gliomas. *Sci. Rep.* **13**, 10389 (2023).

3. M. Venkatesh, J. Jaja, L. Pessoa, Comparing functional connectivity matrices: A geometry-aware approach applied to participant identification. *Neuroimage* **207**, 116398 (2020).

4. D. Vidaurre, A new model for simultaneous dimensionality reduction and time-varying functional connectivity estimation. *PLoS Comput. Biol.* **17**, e1008580 (2021).

5. A. Zalesky, A. Fornito, E. T. Bullmore, Network-based statistic: identifying differences in brain networks. *Neuroimage* **53**, 1197–1207 (2010).

6. D. Vidaurre, A. Llera, S. M. Smith, M. W. Woolrich, Behavioural relevance of spontaneous, transient brain network interactions in fMRI. *Neuroimage* **229**, 117713 (2021).

7. D. S. Margulies, *et al.*, Situating the default-mode network along a principal gradient of macroscale cortical organization. *Proc. Natl. Acad. Sci. U. S. A.* **113**, 12574–12579 (2016).

8. M. Tamietto, *et al.*, Innate cortical gradients constrain cross-modal plasticity. *Research Square* (2025).

9. H. Song, W. M. Shim, M. D. Rosenberg, Large-scale neural dynamics in a shared low-dimensional state space reflect cognitive and attentional dynamics. *Elife* **12** (2023).
